# Gre factors protect against phenotypic diversification and cheating in *Escherichia coli* populations under toxic metabolite stress

**DOI:** 10.1101/2023.01.02.522506

**Authors:** Darshan M. Sivaloganathan, Xuanqing Wan, Mark P. Brynildsen

**Author notes:** **Correspondence:** Mark P. Brynildsen.

## Abstract

Nitric oxide (·NO) is one of the toxic metabolites that bacteria can be exposed to within phagosomes. Gre factors, which are also known as transcript cleavage factors or transcription elongation factors, relieve back-tracked transcription elongation complexes by cleaving nascent RNAs, which allows transcription to resume after stalling. Here we discovered that loss of both Gre factors in *E. coli*, GreA and GreB, significantly compromised ·NO detoxification through a phenotypic diversification of the population. Under normal culturing conditions, both wild-type and Δ*greA*Δ*greB* synthesized protein uniformly. However, treatment with ·NO led to bimodal protein expression in Δ*greA*Δ*greB*, whereas wild-type remained unimodal. Interestingly, exposure to another toxic metabolite of phagosomes, hydrogen peroxide (H_2_O_2_), produced similar results. We found that the diversification in Δ*greA*Δ*greB* cultures required *E. coli* RNAP, occurred at the level of transcription, and could produce cheating where transcriptionally-deficient cells benefit from the detoxification activities of the transcriptionally-proficient subpopulation. Collectively, these results indicate that Gre factors bolster bacterial defenses by preventing phenotypic diversification and cheating in environments with fast-diffusing toxic metabolites.

**Importance:** Toxic metabolite stress occurs in a broad range of contexts that are important to human health, microbial ecology, and biotechnology; whereas Gre factors are highly conserved throughout the bacterial kingdom. Here we discovered that the Gre factors of *E. coli* prevent phenotypic diversification under toxic metabolite stress. Such conformist regulation improves populationwide removal of those stressors and protects against cheating, where one subpopulation commits resources to counter a threat, and the other subpopulation does not, yet both subpopulations benefit.

## Introduction

Phagocytes are instrumental in controlling and eliminating infections as they represent the first line of defense mounted against invading microorganisms [1–4]. Within phagosomes, bacteria are exposed to a wide array of antimicrobials that work in concert to eliminate them [1,2]. One important antimicrobial produced in these environments is nitric oxide (·NO) [2,5–7]. Many microbes have evolved defense systems to combat the cytotoxic effects of ·NO, and the ability to detoxify ·NO has been linked to the virulence of numerous pathogens [8,9]. For example, *S. typhimurium* strains lacking the ·NO dioxygenase, Hmp, displayed reduced survival in mouse models but displayed similar survival to wild-type cells in mice lacking ·NO synthase [10]. Similarly, loss of the ·NO reductase in enterohemorrhagic *E. coli*, through mutation of *norV*, led to a dramatic reduction in survival within human macrophages [11]. Many other bacteria, including *Pseudomonas aeruginosa, Vibrio cholerae*, and *Mycobacterium tuberculosis*, have been shown to rely on ·NO detoxification to impart their virulence [12–14]. Sensitizing pathogens to ·NO by interfering with their defense mechanisms has been suggested to be an appealing approach to antimicrobial therapies, which would be orthogonal to current treatments [8,9,13]. A deeper understanding of the networks that regulate ·NO metabolism will facilitate the discovery of novel, accessible targets that can interfere with the virulence of diverse pathogens [8,9].

Previous work has revealed the importance of the global transcriptional regulator, DksA, to *Salmonella* survival both in macrophages and mouse models [15–17]. Further, DksA has been shown to play an essential role in the ability of *E. coli* to detoxify ·NO with Hmp [18]. DksA is known to bind directly to the secondary channel of RNA polymerase (RNAP) to exert its regulatory functions [19,20]. Interestingly, DksA is member of a larger classes of proteins that collectively compete for access to the secondary channel of RNAP to exert their functions [21– 25]. Additional secondary channel proteins include Gre factors (also known as transcript cleavage factors or transcription elongation factors), the nucleoside diphosphate kinase regulator Rnk, the conjugative F plasmid regulator TraR, and the antimicrobial peptide microcin J25 [26– 29]. While recent research has solidified a central role for DksA in bacterial ·NO defenses, the impacts of other RNAP secondary channel regulators on bacterial ·NO responses remain ill-defined.

Gre factors are highly conserved proteins across the bacterial kingdom that alleviate RNAP stalling and promote transcript elongation [27,30,31]. RNAP stalling is thought to occur as a result of numerous events including abortive initiation, misincorporation of nucleotides, and collisions with DNA-bound protein complexes [32]. RNAP stalling often leads to reverse translocation, where the transcription elongation complex (TEC) slides backwards in the 3’ to 5’ direction along the nascent RNA transcript, exposing one or more base pairs [32,33]. Backtracked TECs cannot proceed with transcript elongation, and when held in a backtracked position for prolonged periods of time, they can eventually lead to an arrested transcriptional state, and ultimately dissociation of RNAP [32,33]. Gre factors counteract RNAP stalling by facilitating the 3’ cleavage of nascent RNA in backtracked TECs, allowing transcription to resume [30,32]. The importance of Gre factors to pathogenicity has been established for a number of pathogens [34–38], where it has been shown to regulate a diversity of virulence factors [35,36].

Inspired by previous work on DksA and ·NO stress, the involvement of Gre factors in the pathogenicity of various microbes, and the fact that both act through the secondary channel of RNAP, we investigated the importance of GreA and GreB to ·NO detoxification by *E. coli*. We found that ·NO detoxification was robust to individual deletion of GreA or GreB; however, the double mutant, Δ*greA*Δ*greB*, was significantly impaired. Further analyses revealed that loss of the Gre factors led to phenotypic diversification under ·NO stress, where a subpopulation produced Hmp, the main aerobic ·NO detoxification enzyme, similarly to wild-type and another subpopulation failed to produce Hmp appreciably. Diversification was observed with an orthogonal expression system under ·NO stress, whereas distributions were unimodal under normal growth conditions. Experiments with fluorescently labeled RNA, fluorescent proteins, and an independent T7 RNAP expression system established that diversification in RNA synthesis was a major determinant of bimodal protein distributions and that both depended on *E. coli* RNAP. Importantly, ·NO diffuses rapidly and is catalytically detoxified, which suggested that Hmp-deficient subpopulations may cheat off their Hmp-proficient neighbors, which commit resources to deal with the threat. Interestingly, using a model system of mixed populations consisting of wild-type and Δ*hmp*, we confirmed that the phenotypic diversification observed in Δ*greA*Δ*greB* cultures would generate cheating dynamics. In addition, we found that the phenotypic diversification of Δ*greA*Δ*greB* populations is not confined to ·NO stress, but is also observed in the presence of hydrogen peroxide (H_2_O_2_), which is another phagosome antimicrobial. Overall, these results suggest that the Gre factors prevent phenotypic diversification and cheating in *E. coli* populations under toxic metabolite stress, and contribute to the growing understanding of how regulators that bind to the secondary of channel of RNAP modulate bacterial defense networks.

## Results

### 2.1 Loss of GreA and GreB impairs ·NO detoxification

To begin to explore the role of Gre factors in ·NO defenses, we constructed single and double deletion mutants of GreA and GreB, and measured their abilities to detoxify ·NO compared to wild-type (WT). Aerobic cultures were grown to mid-exponential phase and treated with 250 μM of the ·NO donor, DPTA NONOate (Fig 1A). For all cultures, ·NO concentrations rapidly rose and peaked around 4 μM. The clearance times, which we define as the time it takes for [·NO] to fall below 0.5 μM, were approximately 0.4 hr for WT, Δ*greA*, and Δ*greB*. However, for Δ*greA*Δ*greB*, the time to eliminate ·NO from cultures was significantly longer at approximately 1.4 hr, representing a greater than 3-fold delay in detoxification. Additionally, Δ*greA*Δ*greB* cultures had impaired growth resumption following ·NO stress compared to WT and single deletion mutants (Fig 1B). Using genetic complementation experiments, we confirmed that the observed phenotype was attributable to loss of both GreA and GreB. Introduction of either *greA* or *greB* when expressed from their native promoters on low-copy plasmids restored ·NO detoxification to WT levels (Fig S1A). We also observed indistinguishable culturability levels between Δ*greA*Δ*greB* and WT under ·NO stress (Fig S1B), which suggested that cell death was not a contributing factor to the impaired ·NO detoxification observed.

**Figure 1.**
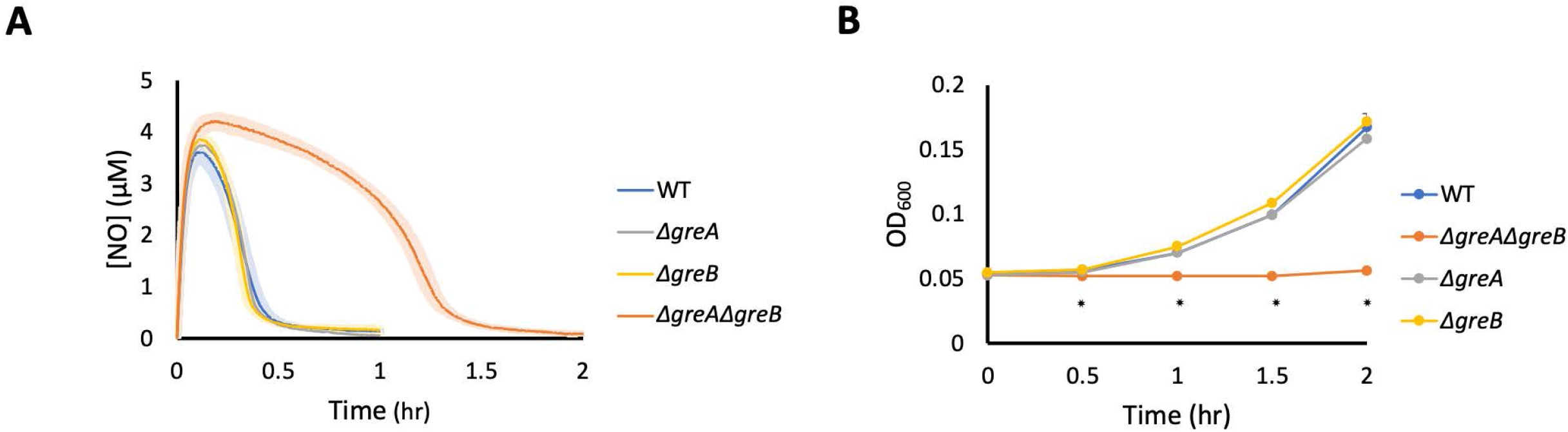
Loss of GreA and GreB significantly impaired ·NO detoxification and growth resumption. Cultures were grown in MOPS minimal media to mid-exponential phase and inoculated into a bioreactor at an OD600 of 0.05. Immediately after, 250 μM of DPTA NONOate was added to the bioreactor. WT (blue), ΔgreA (grey), ΔgreB (yellow), ΔgreAΔgreB (orange). (A) ·NO concentrations were continuously monitored in the bioreactor. Solid lines represent the mean of at least three independent replicates, whereas light shading represents the standard error of the means. (B) Samples were removed to measure OD600 at indicated time points. Circles represent the mean of at least three independent replicates, whereas error bars represents the standard error of the means. Asterisks indicate statistical significance, which was assessed by one-way ANOVA.

### 2.2 Δ*greA*Δ*greB* do not display a growth defect in the absence of ·NO stress

Given the impairment in ·NO detoxification, we considered whether Δ*greA*Δ*greB* in general suffered from growth or translational defects, since the main determinant of ·NO clearance time in aerobic cultures is de novo synthesis of Hmp [18,39,40]. To investigate this, we compared the growth of both WT and Δ*greA*Δ*greB* cultures harboring an IPTG-inducible, GFP reporter plasmid. Notably, Δ*greA*Δ*greB* cultures grew significantly faster from lag to mid-exponential phase than WT in the minimal media used here (Fig 2A), and GFP was produced to significantly higher levels in Δ*greA*Δ*greB* than WT (Fig 2B, S1C & S1D). The growth advantage depicted in Figure 2 was not dependent on the GFP reporter plasmid (Fig S1E), and GFP fluorescence was negligible in the absence of the GFP reporter plasmid and IPTG (Fig S1F). These results demonstrated that during normal, non-stressed growth in minimal media Δ*greA*Δ*greB* did not exhibit growth defects or impairments in protein synthesis, which suggested that impairments in ·NO detoxification did not originate from general growth or translational issues with the mutant.

**Figure 2.**
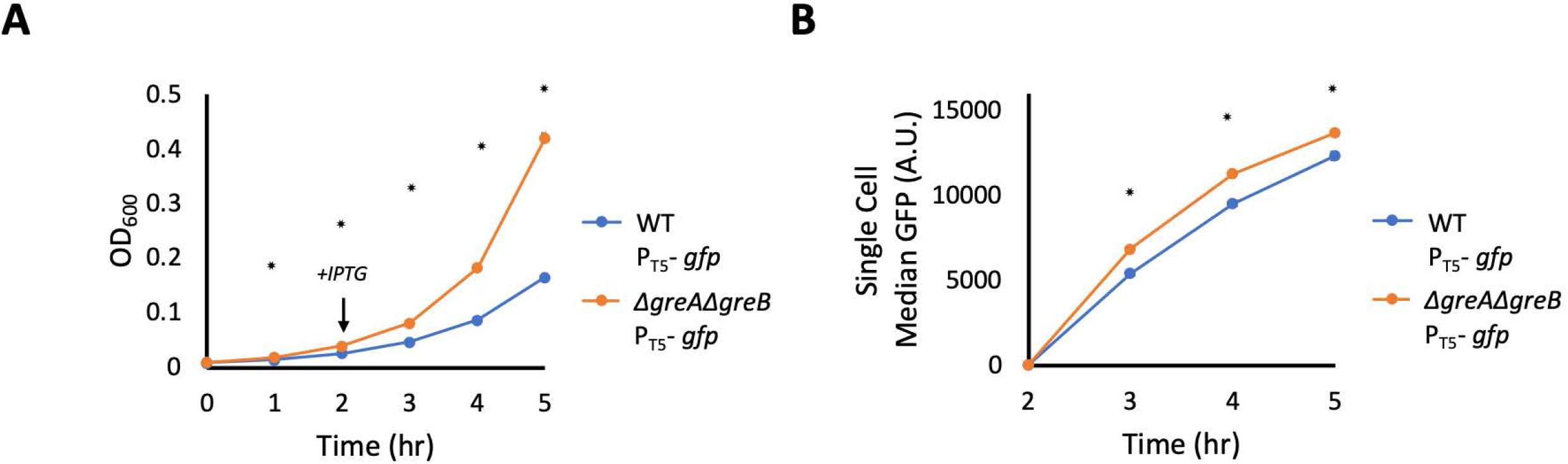
Δ*greA*Δ*greB* display growth and protein production advantages in the absence of ·NO stress. WT and Δ*greA*Δ*greB* cultures were inoculated into baffled flasks containing MOPS minimal media at and OD_600_ of 0.01 and incubated at 37 °C and 250 RPM. (A) Samples were removed to measure OD_600_ at indicated time points. (B) After 2 hrs of incubation, 1 mM IPTG was added to flasks to induce GFP production. Samples were removed, fixed, and median GFP fluorescence was obtained by flow cytometry. WT (blue), Δ*greA*Δ*greB* (orange). Circles represent the means of at least 3 independent replicates. Error bars represent the standard errors of the means. Asterisks indicate statistical significance, at a p-value ≤ 0.05, which was assessed by t-tests.

### 2.3 Δ*greA*Δ*greB* exposed to ·NO displayed bimodal protein expression from the *hmp* promoter

The delayed ·NO detoxification of Δ*greA*Δ*greB* suggested that cells lacking Gre factors have impaired translation under ·NO stress. In *E. coli*, the enzyme Hmp detoxifies the majority of ·NO under aerobic conditions, and previous work has established that it is largely synthesized de novo in response to ·NO [18,39,41]. To confirm that similar circumstances govern ·NO detoxification under the conditions used here, we deleted *hmp* from WT and Δ*greA*Δ*greB* and measured ·NO consumption. Notably, Δ*hmp* and Δ*greA*Δ*greB*Δ*hmp* cultures both failed to detoxify NO appreciably (Fig S1G), which suggested that the difference observed between the strains required *hmp*. To assess the importance of protein production under ·NO stress, we treated both strains with 100 μg/mL chloramphenicol (CAM) prior to and during ·NO treatment, and observed negligible ·NO detoxification in WT and Δ*greA*Δ*greB* cultures (Fig S1H). These results established that the difference in ·NO detoxification between WT and Δ*greA*Δ*greB* required translation under NO stress.

Since translation and *hmp* were required to observe differences in ·NO detoxification between WT and Δ*greA*Δ*greB*, we used a transcriptional reporter of *hmp* expression (P_hmp_-gfp) in Δ*hmp* and Δ*greA*Δ*greB*Δ*hmp* to monitor protein production from that promoter during ·NO stress. We opted to use strains devoid of *hmp* here to ensure identical ·NO environments during the assay (Fig S1G). Interestingly, we observed that Δ*hmp* and Δ*greA*Δ*greB*Δ*hmp* displayed drastically different fluorescence distributions (Fig 3). Populations of Δ*hmp* expressed GFP in a unimodal fashion that increased in magnitude over time. In contrast, Δ*greA*Δ*greB*Δ*hmp* populations expressed GFP in a bimodal fashion, in which one subpopulation of cells did not increase expression of GFP over time, whereas the other subpopulation expressed GFP at levels similar to Δ*hmp*. Complementation experiments with expression of either *greA* or *greB* from their native promoters restored unimodal GFP distributions (Fig S2). These results suggested that the impairment in ·NO detoxification in Δ*greA*Δ*greB* originated from bimodal Hmp expression, where a sizable subpopulation of cells did not express Hmp in response to ·NO.

**Figure 3.**
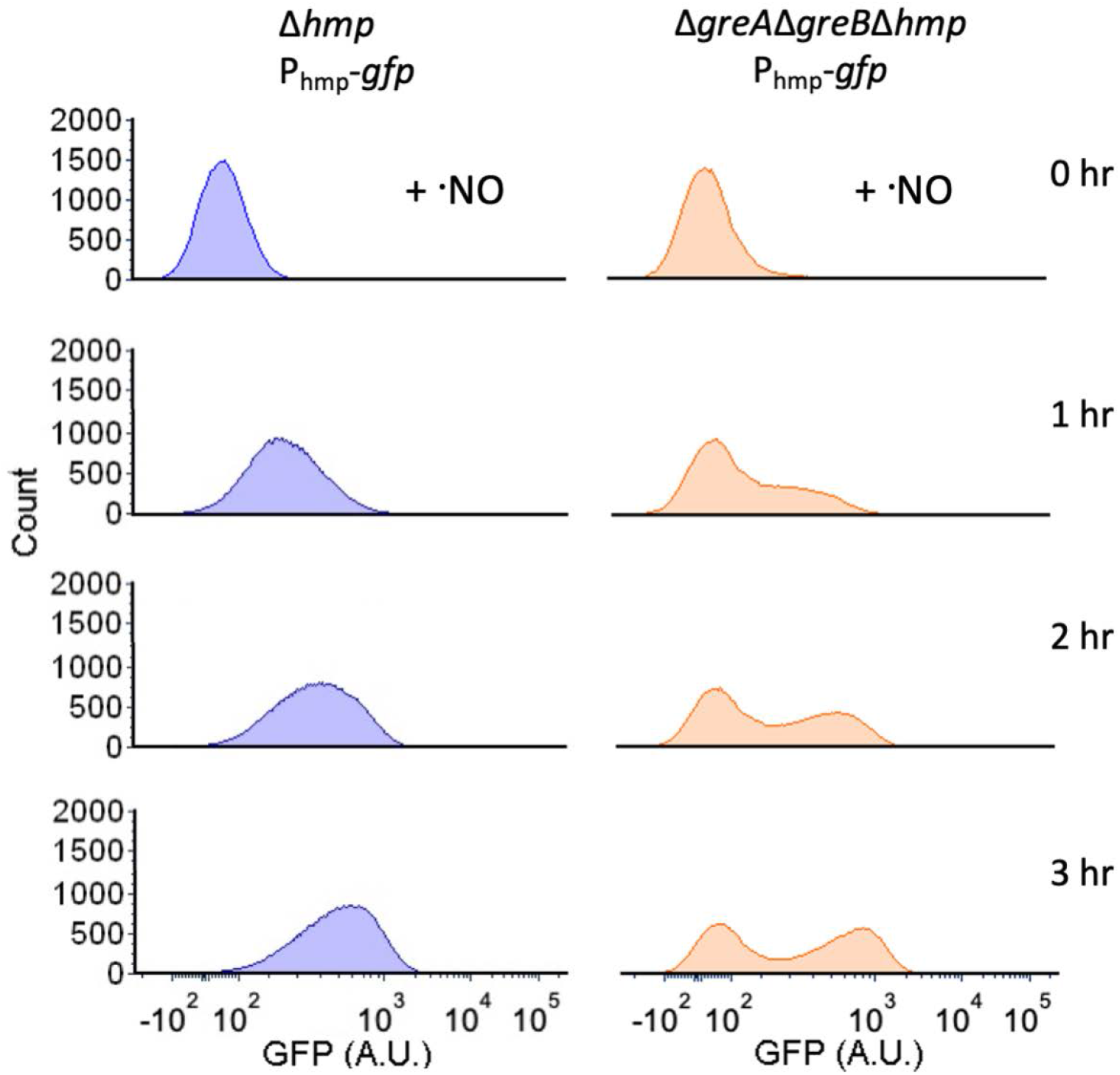
Under ·NO stress, Δ*greA*Δ*greB* populations exhibit bimodal protein expression from the *hmp* promoter. Cultures were grown in MOPS minimal media to mid-exponential phase and inoculated into a bioreactor at an OD_600_ of 0.05. Immediately after, 250 μM of DPTA NONOate was added. Samples were removed at indicated time points, fixed, and GFP fluorescence was measured by flow cytometry. Histograms represent single cell GFP fluorescence distributions at indicated time points. Δ*hmp* (blue) and Δ*greA*Δ*greB*Δ*hmp* (orange). Images are representative of at least 3 biological replicates.

### 2.4 Expression bimodality extends to an orthogonal expression system

We were interested to see whether the bimodality observed from the *hmp* promoter extended to other expression systems. To test this, we expressed GFP from an IPTG-inducible P_T5_ promoter under ·NO stress. We found that expression was again bimodal with this independent expression system (Fig 4A). Using an Hmp-GFP translational fusion in place of GFP in the same expression construct we observed delayed ·NO detoxification (Fig 4B), and bimodal Hmp-GFP expression in Δ*greA*Δ*greB*Δ*hmp*, which was absent from Δ*hmp* (Fig 4C). Collectively, these data suggested that bimodal protein expression in Δ*greA*Δ*greB* under ·NO stress was not specific to the *hmp* promoter, but rather it reflected a more general trait of the Gre factor mutant. Moreover, use of Hmp-GFP provided direct evidence of an association between impaired ·NO detoxification and bimodal protein expression in Δ*greA*Δ*greB* under ·NO stress.

**Figure 4.**
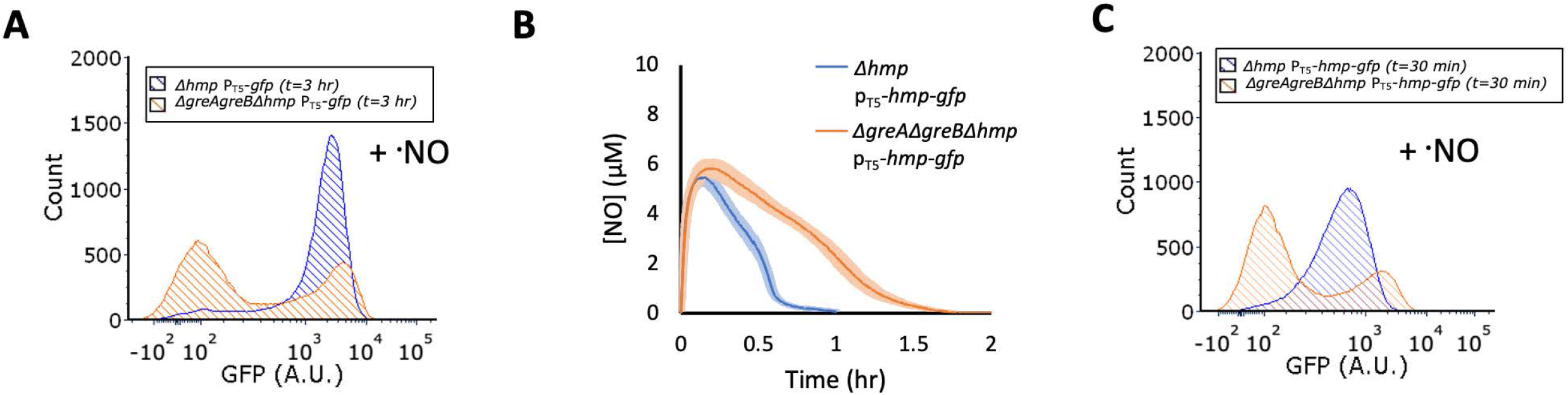
Bimodal protein expression in Δ*greA*Δ*greB* populations extends to the T_5_ promoter. Cultures were grown in MOPS minimal media to mid-exponential phase and inoculated into bioreactors at an OD_600_ of 0.05. Immediately after, 250 μM (Panel A) or 350 μM (Panels B and C) of DPTA NONOate were added, along with 1 mM IPTG. A slightly higher DPTA NONOate concentration was used for experiments with Hmp-GFP to match peak ·NO concentrations from other experiments. Samples were removed at indicated time points, fixed, and fluorescence was measured by flow cytometry. Panel (A) reflects fluorescence distributions for cells expressing GFP under the T_5_ promoter. Panels (B) and (C) reflect ·NO concentration profiles, and fluorescence distributions, respectively, for cells expressing Hmp-GFP translational fusions under the T_5_ promoter. ·NO curves (Panel B) show the means of three independent replicates and error bars represent the standard errors of the means. Flow cytometry with Hmp-GFP are depicted at an earlier time point (30 min), such that ·NO would still be present in both cultures. Δ*hmp* (blue) and Δ*greA*Δ*greB*Δ*hmp* (orange). Fluorescent histograms are representative of three or more biological replicates. Fluorescent histograms at t=0 are displayed in Figure S3.

### 2.5 Bimodal protein expression arises from a phenotypic diversification

The observation that two populations arise under ·NO stress led us to question whether the phenomenon was heritable, and thus originating from a genetic change, or whether it was transient, and originating from a phenotypic diversification. To answer this question, we exposed Δ*greA*Δ*greB*Δ*hmp* cultures to ·NO and sorted cells from the top 10% (high) and bottom 10% (low) of the fluorescence distribution after 1 hr of ·NO exposure (Fig 5). Those subpopulations were then cultured and prepared for subsequent assays. Interestingly, cells originating from the high and low subpopulations displayed the same bimodal GFP expression pattern as the parent population after outgrowth, which demonstrated that the fluorescence/protein expression status of subpopulations was not heritable, but transient. Cell culturability was similar between populations (Fig S4A & S4B), and neither subpopulation had a significant difference in terminal cell density or growth dynamics (Fig S4C & S4D). These data showed that bimodal protein expression in ·NO-stressed, Δ*greA*Δ*greB* populations arose from phenotypic diversification, rather than genetic mutation.

**Figure 5.**
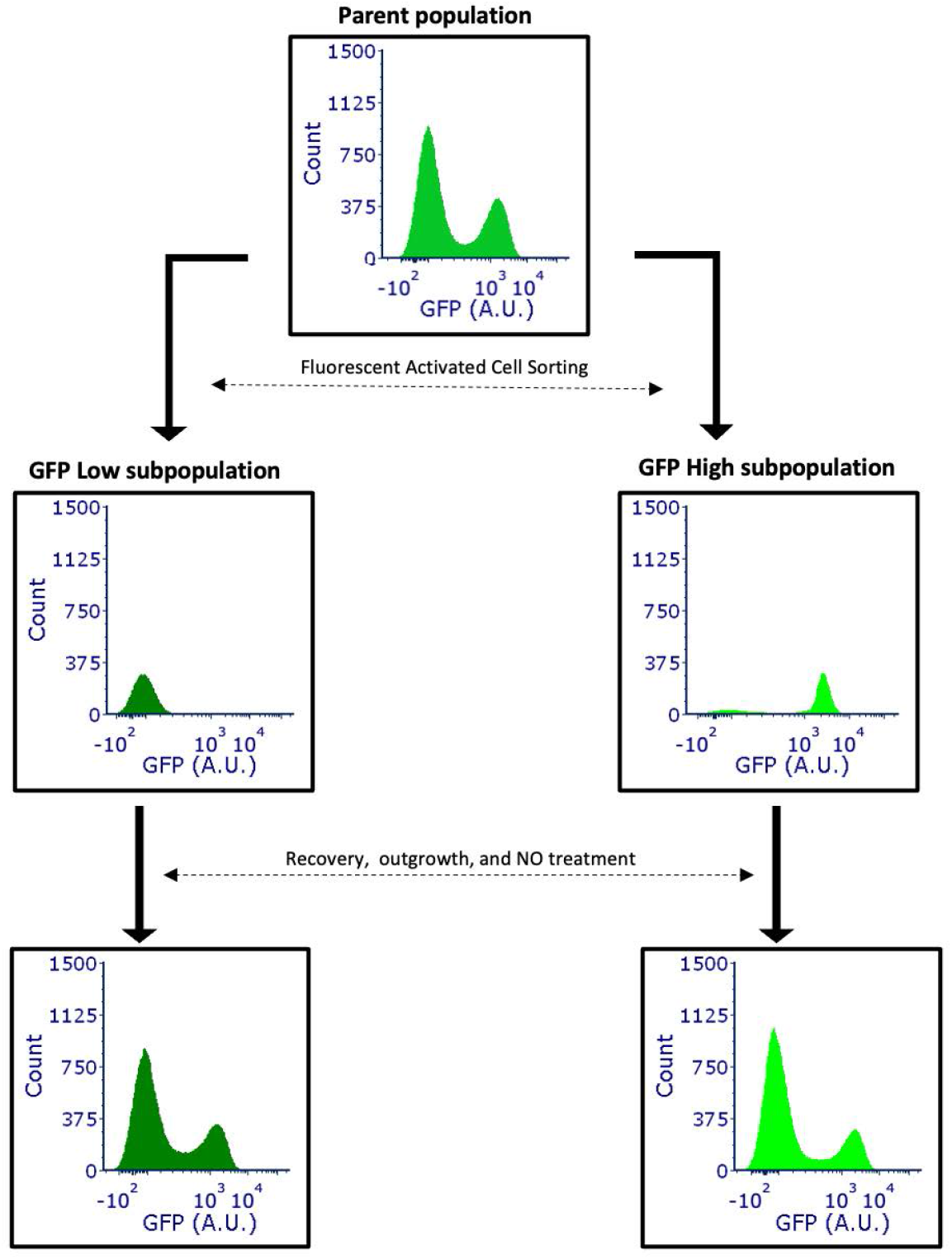
Δ*greA*Δ*greB*Δ*hmp* cells do not retain memory of GFP expression under prior NO stress. Cultures were grown in MOPS minimal media to mid-exponential phase and inoculated into bioreactors at an OD_600_ of 0.05. Immediately after, 250 μM of DPTA NONOate and 1 mM IPTG were added to the bioreactor. Samples were removed at 1 hr and passed through a cell sorter to collect 1 million cells from the bottom 10 percent and top 10 percent of the fluorescence distribution. Collected cells were recovered in rich media, grown to mid-exponential phase in MOPS minimal media, and dosed with 250 μM DPTA NONOate and 1 mM IPTG. Samples were removed after 1 hr, fixed and fluorescence distributions were measured by flow cytometry. Parent population (medium shade green), GFP low subpopulation (dark shade green), GFP high subpopulation (light shade green).

### 2.6 Phenotypic diversification at the level of transcription

To further understand this phenomenon, we sought to assess whether the phenotypic diversification we observed at the protein level was present at the level of transcription or whether it originated from a post-transcriptional process. To accomplish this, we used an RNA aptamer (tdBroccoli) capable of emitting green fluorescence upon complexing with a cell permeable, non-fluorescent fluorophore (DFHBI-1T) (Fig S5A) [42–45]. Neither tdBroccoli, nor DFHBI-1T produced appreciable fluorescence on their own (Fig S5B) [44]. We expressed tdBroccoli from an IPTG-inducible promoter in Δ*hmp* and Δ*greA*Δ*greB*Δ*hmp* cultures. In the absence of ·NO, both Δ*hmp* and Δ*greA*Δ*greB*Δ*hmp* cells displayed unimodal fluorescence distributions (Fig S5C). However, in the presence of ·NO stress, we found that Δ*greA*Δ*greB*Δ*hmp* cells displayed bimodal fluorescence, suggesting that phenotypic diversification at the protein level included contributions from diversification at the level of transcription (Fig 6). To demonstrate this directly, we transcriptionally fused mCherry to the 5’ end of tdBroccoli [45,46]. Such an expression system allows direct quantification of both transcriptional and protein output, simultaneously. We found that, under ·NO stress, Δ*hmp* cells displayed a unimodal expression pattern in both transcript and protein (Fig 7A-B) with the vast majority of cells increasing transcript and protein levels concurrently (Fig 7C). On the other hand, Δ*greA*Δ*greB*Δ*hmp* displayed bimodal transcript and protein expression patterns (Fig 7D-E), where one subpopulation simultaneously contained low transcript and protein abundance, and the other subpopulation concurrently increased transcript and protein expression (Fig 7F). Collectively, these data suggested that, under ·NO stress, Δ*greA*Δ*greB* cells diversify into two subpopulations, one inactive subpopulation impaired in both transcript and protein synthesis and a second active subpopulation producing transcript and protein at levels comparable to WT.

**Figure 6.**
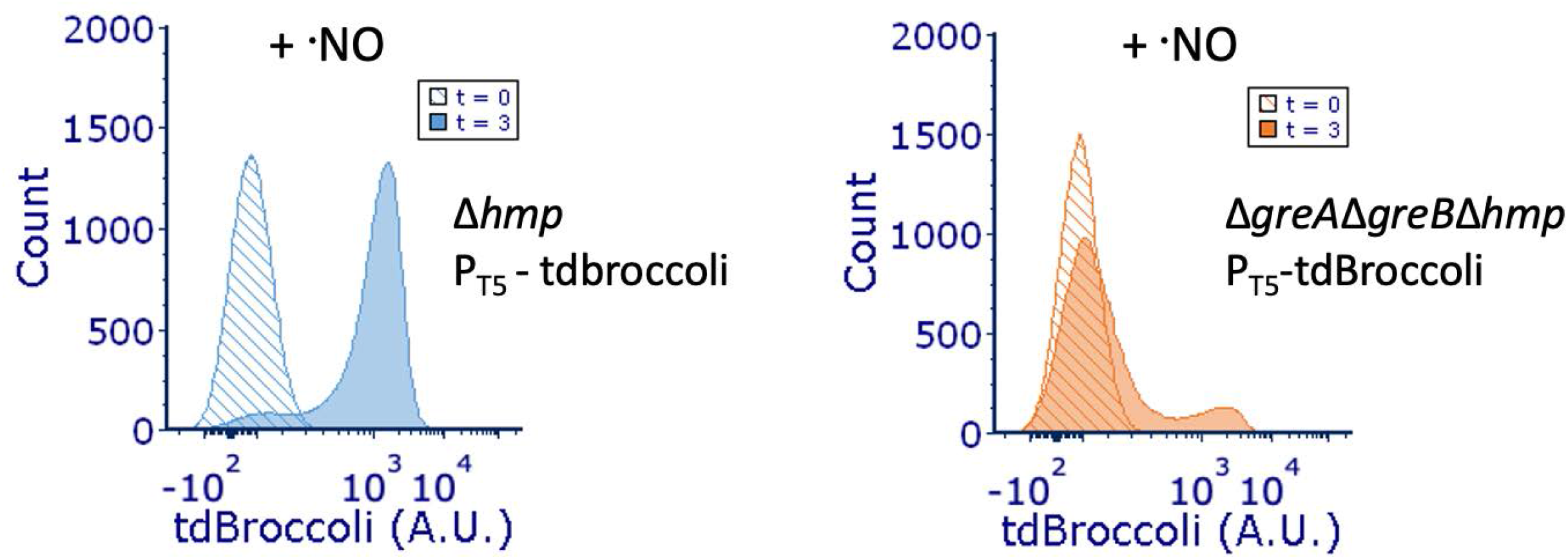
Δ*greA*Δ*greB* display bimodal transcript production. Cultures of Δ*hmp* and Δ*greA*Δ*greB*Δ*hmp* harboring P_T5_-tdbroccoli were grown in MOPS minimal media to midexponential phase and inoculated into a bioreactor at an OD_600_ of 0.05. Immediately after, 250 μM of DPTA NONOate, 1 mM IPTG and 50 μM DFHBI-1T were added to the bioreactor. Fluorescence distributions at the 0 and 3-hr time points are depicted. Δ*hmp* (blue) and Δ*greA*Δ*greB*Δ*hmp* (orange). Images are representative of at least three biological replicates.

**Figure 7.**
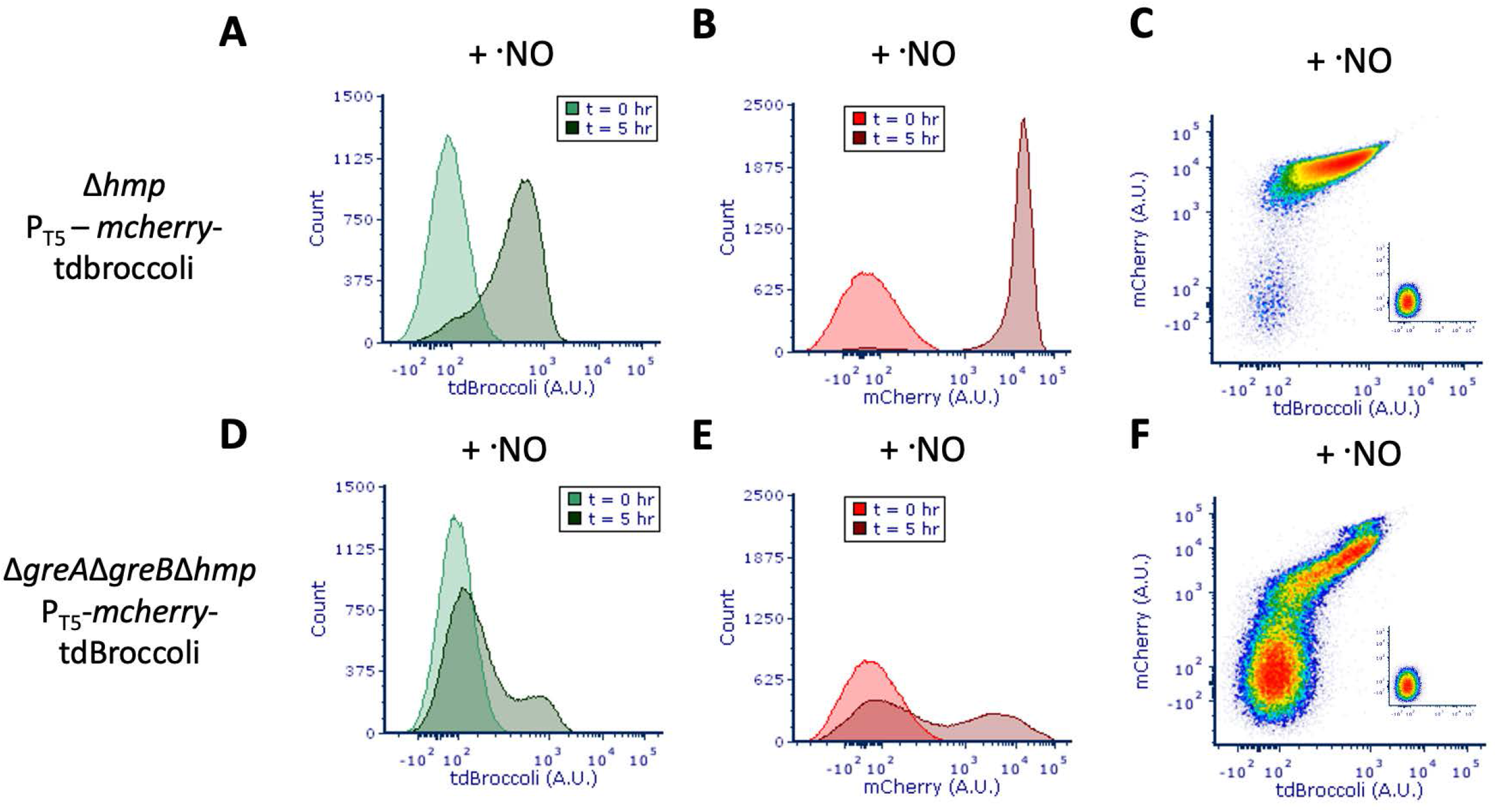
Positive correlation between transcript and protein expression in Δ*greA*Δ*greB* subpopulations. Cultures of Δ*hmp* and Δ*greA*Δ*greB*Δ*hmp* harboring P_T5_-*mcherry*-tdbroccoli were grown in MOPS minimal media to mid-exponential phase and inoculated into a bioreactor at an OD_600_ of 0.05. Immediately after, 250 μM of DPTA NONOate, 2 mM IPTG and 50 μM DFHBI-1T were added to the bioreactor. Five hrs after treatment, samples were removed and tdBroccoli and mCherry fluorescence were measured concurrently by flow cytometry. Panels (A) and (B) represent Δ*hmp* tdBroccoli and mCherry, distributions respectively. Panels (D) and (E) depict Δ*greA*Δ*greB*Δ*hmp* tdBroccoli and mCherry, respectively. Panels (C) and (F) represent tdBroccoli vs. mCherry distributions for Δ*hmp* and Δ*greA*Δ*greB*Δ*hmp* respectively at t = 5 hr, whereas subplots represent t = 0 hr. Images are representative of at least three biological replicates.

### 2.7 Phenotypic diversification can solely be attributed to transcription

We sought to determine whether post-transcriptional events also contributed to phenotypic diversification. To assess this, we used the bacteriophage T_7_ RNA polymerase (T_7_ RNAP), a non-native, single unit polymerase, that is evolutionarily distinct from *E. coli* RNAP, and is not known to interact with Gre factors [47,48]. T_7_ RNAP was expressed from a constitutive *E. coli* promoter, so that it would be present as a functional protein during assays, and rifampicin was added to block further transcription from *E. coli* RNAP (including that of T_7_ RNAP), whereas *gfp* was expressed from an IPTG-inducible T_7_ promoter, which are not recognized by *E. coli* RNAP (Fig S6). With this design, transcription of *gfp* under rifampicin treatment would be solely attributable to T_7_ RNAP, without a contribution from *E. coli* RNAP, and thus post-transcriptional events could be assessed for phenotypic diversification independent of *E. coli* RNAP. Using this system with Δ*hmp* and Δ*greA*Δ*greB*Δ*hmp*, cells were treated with ·NO and rifampicin, and IPTG was added to induce GFP production. We observed a unimodal distribution pattern for both Δ*hmp* and Δ*greA*Δ*greB*Δ*hmp* cultures expressing GFP from the T_7_ promoter (Fig 8). Switching to a non-native RNAP expression system eliminated bimodal protein expression, which provided evidence that post-transcriptional mechanisms and NTP depletion did not contribute to the phenomenon. Further, it suggested that phenotypic diversification was dependent upon transcription performed by *E. coli* RNAP.

**Figure 8.**
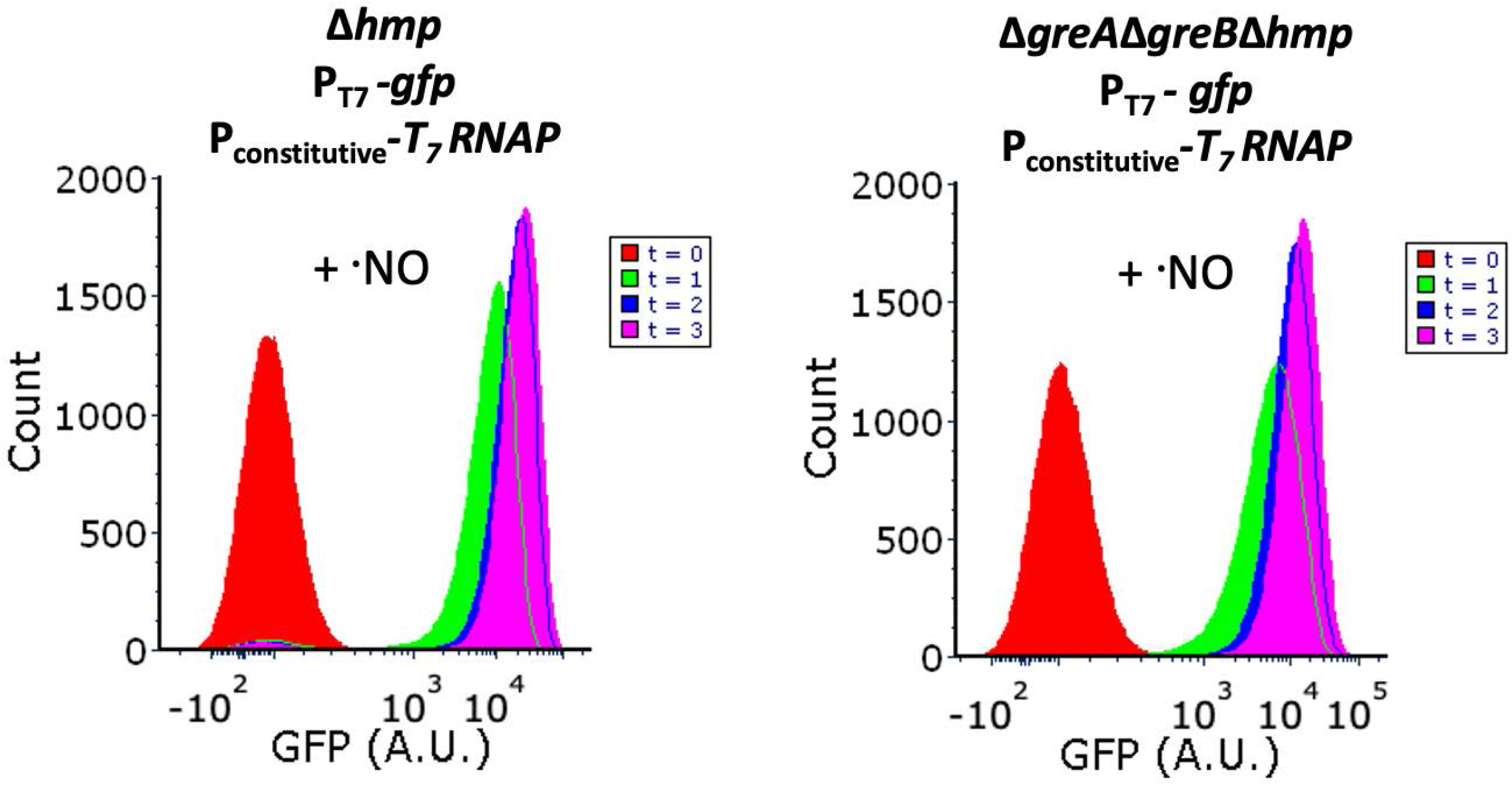
GFP expression from orthogonal transcription system produces unimodal protein expression in Δ*greA*Δ*greB* populations. Δ*hmp* and Δ*greA*Δ*greB*Δ*hmp* cultures harboring T_7_ RNAP under a constitutive promoter (J23114) and P_T7_-*gfp* were grown in MOPS minimal media to mid-exponential phase and inoculated into bioreactors at an OD_600_ of 0.05. Immediately after, 100 μg/mL rifampicin was added to the bioreactor. After 10 min, 250 μM of DPTA NONOate and 1 mM IPTG were added. Samples were removed at t = 0, 1, 2 and 3 hr, fixed and GFP fluorescence distributions were obtained by flow cytometry analysis. Images are representative of at least 3 biological replicates.

### 2.8 Gre factors prevent cheating under ·NO stress

Our results suggested that loss of Gre factors produced phenotypic diversification under ·NO stress, which manifested as two subpopulations with significantly different levels of the detoxification enzyme Hmp. Previous work established that Hmp detoxifies approximately 99.8% of intracellular ·NO under similar conditions, and that after a short induction period cellular ·NO detoxification vastly exceeds abiotic loss of ·NO *(e.g*., autooxidation, gas phase transport) [39]. In consideration of this knowledge and the fact that ·NO is a small uncharged molecule that diffuses rapidly, we reasoned that the burden of ·NO detoxification in Δ*greA*Δ*greB* cultures would fall upon the Hmp-expressing subpopulation, whereas the non-responsive subpopulation would not need to commit resources to deal with ·NO, but could benefit from the other subpopulation clearing ·NO from the environment. Such a scenario would constitute cheating by the non-responsive subpopulation, which is not present in WT cultures. To examine the plausibility of this scenario, we performed experiments with monocultures and cocultures of Hmp-proficient and Hmp-deficient strains. To enable differentiation of strains in coculture, we labeled them with different fluorophores. Specifically, the Hmp-proficient strain constitutively expressed mCherry, whereas the Hmp-deficient strain constitutively expressed GFP. Consistent with previous findings, monocultures of the Hmp-proficient strain cleared ·NO and resumed growth significantly faster than monocultures of Hmp-deficient strains (Fig 9). Further, we found that 50/50 cocultures of Hmp-proficient and Hmp-deficient strains cleared ·NO slower than the Hmp-proficient monoculture (Fig 9A), and when growth was monitored, the fold-change in cell number for the Hmp-proficient and Hmp-deficient strains in coculture were comparable and both lower than the Hmp-proficient monoculture (Fig 9B). These data suggested that the phenotypic diversification observed in Δ*greA*Δ*greB* in response to ·NO produced a scenario where one subpopulation could cheat on the altruistic behavior of another subpopulation.

**Figure 9.**
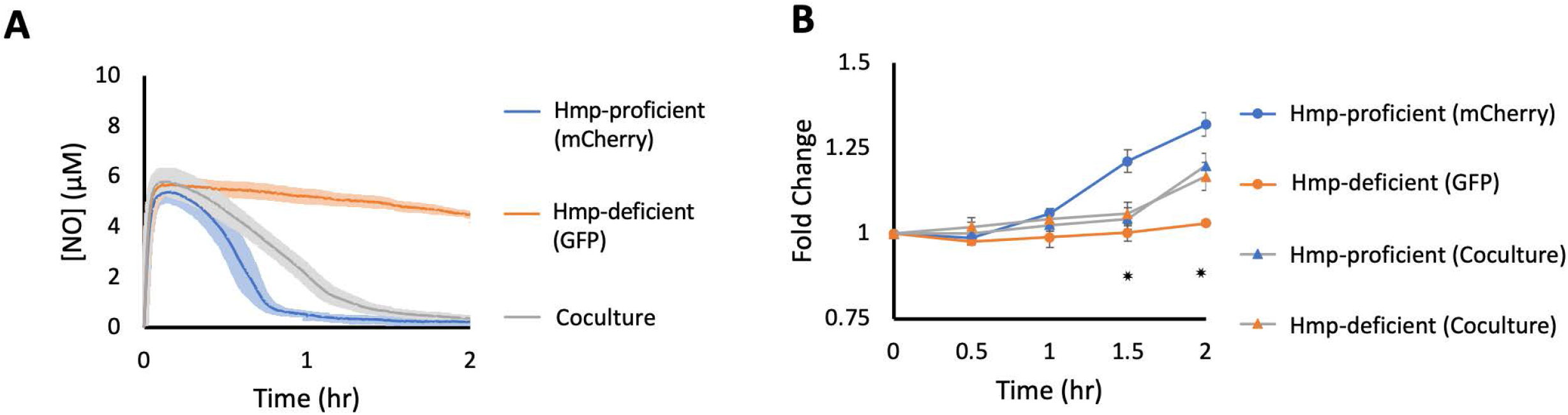
Cheating occurs in ·NO-treated cocultures of Hmp-proficient and Hmp-deficient strains. Cultures were grown in MOPS minimal media to mid-exponential phase and inoculated into a bioreactor at an OD_600_ of 0.025 either as a monoculture of Hmp-proficient cells (blue), a monoculture of Hmp-deficient cells (orange), or as a 1:1 coculture of both strains (grey). Immediately after, 250 μM of DPTA NONOate was added to the bioreactor. (A) ·NO concentrations were continuously monitored in the bioreactor. Solid lines represent the means of at least three independent replicates, whereas light shading represents the standard errors of the means. (B) Samples were removed to measure OD_600_ at indicated time points. For co-cultures, samples were fixed and the OD_600_ was scaled by the proportion of GFP- and mCherry-positive cells measured by flow cytometry. Colored circles and triangles represent the means of at least three replicates, whereas error bars represent the standard errors of the means. Asterisks indicate statistical significance, which was assessed by one-way ANOVA.

### 2.9 Phenotypic diversification without Gre factors under H_2_O_2_ stress

We sought to assess the generality of stress-induced phenotypic diversification of Δ*greA*Δ*greB* populations, and elected to test H_2_O_2_, which is another toxic metabolite present within phagosomes that is small, uncharged, catalytically detoxified by enzymes, and potentially genotoxic depending on its concentration [8,41,49]. To test this, we expressed GFP from an IPTG-inducible P_T5_ promoter in WT and Δ*greA*Δ*greB* in the presence and absence of H_2_O_2_. We found unimodal expression for both WT and Δ*greA*Δ*greB* in the absence of H_2_O_2_, whereas exposure to 75 μM H_2_O_2_ led to bimodal GFP expression in Δ*greA*Δ*greB* populations and unimodal expression in WT (Fig 10). We note that 25 and 50 μM doses were also tested, and while bimodality in 50 μM treatments of Δ*greA*Δ*greB* begins to emerge at 1 h, the 25 μM treatment remained unimodal through 1h (Fig S7A). Previous work in similar conditions established that 25 μM treatments of H_2_O_2_ are cleared by 1 h [49], and thus these data would suggest that the duration of time under stress is impactful to whether bimodal protein distributions have time to develop in Δ*greA*Δ*greB* populations under stress. Collectively, these data show that phenotypic diversification of Δ*greA*Δ*greB* populations is not confined to ·NO stress, but extends to another prevalent toxic metabolite, H_2_O_2_.

**Figure 10.**
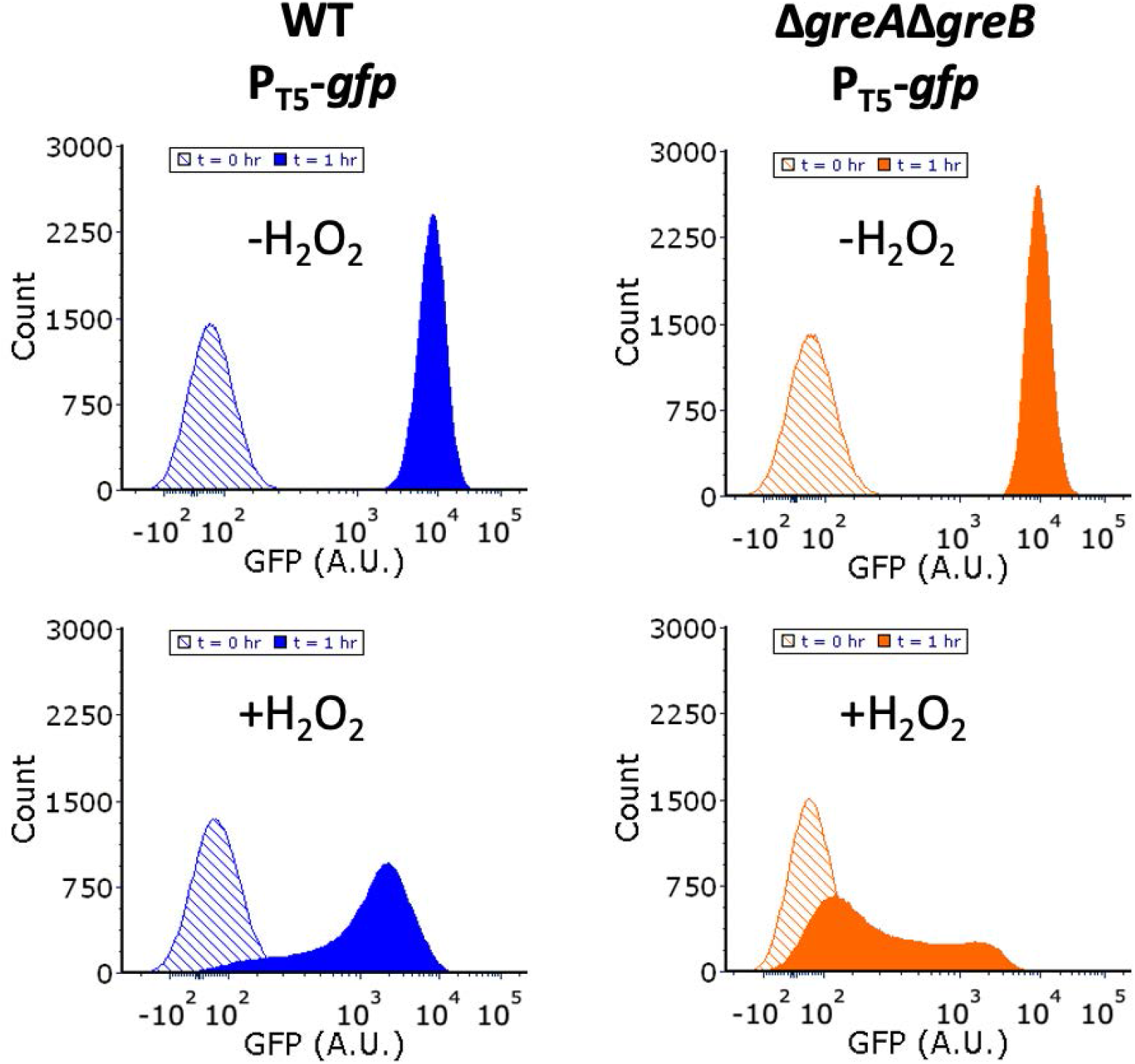
Bimodal protein expression without Gre factors occurs under H_2_O_2_ stress. Cultures were grown in M9 minimal media to mid-exponential phase and inoculated into a bioreactor containing 75 μM H_2_O_2_ (+H_2_O_2_ condition) or the same volume of autoclaved Milli-Q water (-H_2_O_2_ condition). Immediately after 1 mM IPTG was added. Samples were removed at 0 and 1 hr, fixed and GFP distributions were measured by flow cytometry. WT (blue), Δ*greA*Δ*greB* (orange). Images are representative of at least 3 biological replicates.

## Discussion

·NO is a broad-spectrum antimicrobial, and a sizeable number of bacteria rely on ·NO detoxification systems to impart their virulence [8,10,11,50]. As such, increased understanding of bacterial ·NO defense networks could identify novel therapeutic targets for the treatment of bacterial infections [8,9,18,39,40,51–54]. Recently, the global transcriptional regulator, DksA, has been revealed to play an important role in both *S. Typhimurium* ·NO defenses and virulence, as well as ·NO detoxification in *E. coli* [15,16,18]. DksA is a member of a larger class of proteins that exert their function by binding to the secondary channel of RNAP [23,27]. Among these proteins are Gre factors, that share structural homology with DksA [21,22,24]. Given the importance of DksA to the ·NO stress response of *E. coli*, we hypothesized that other regulators of the secondary channel of RNAP may influence ·NO detoxification.

Acting to relieve transcriptional stalling, Gre factors are a widely conserved class of proteins across the prokaryotic kingdom that interact with the secondary channel of RNAP [30,31,33]. Transcriptional stalling is a ubiquitous event, thought to occur during a variety of biochemical activities, including initiation of transcription, recognition of transcriptional errors, collisions with DNA-bound protein complexes, DNA damage, and nucleotide deprivation [21,32,55,56]. As such, Gre factors play a crucial role in transcriptional homeostasis, by aiding in RNAP promoter escape, rescuing arrested RNAP complexes, and even serving a proof-reading role by cleaving mis-incorporated nucleotides from nascent transcripts [21,33]. Gre factors have also been implicated in the survival of numerous microbes in harsh and restrictive environments. For example, loss of GreA in *M. tuberculosis* and *Mycobacterium smegmatis* led to a significant reduction in survival within macrophages [34]. In *Salmonella*, Gre factors were found to be important for pathogenicity and required for expression of genes involved in epithelial invasion, intracellular survival, and replication within hosts cells [35]. Inactivation of GreA in *Francisella tularenensis* led to reduced invasion and growth within macrophages, as well as reduced survival in mouse models [36]. For numerous other bacteria, Gre factors have been implicated as stress proteins, upregulated under harsh environmental conditions that include acid stress, oxidative stress, heat shock, hypoxia, and salt stress [57–61].

The evidence suggesting that Gre factors play an important role in intracellular pathogen survival, coupled with their similarities to DksA, inspired us to explore whether such factors play a role in modulating the ·NO stress response of *E. coli*. We found that individual deletion of either *greA* or *greB* had no impact; however, combined loss of both genes significantly impaired ·NO detoxification (Fig 1). Cells lacking *greA* and *greB* did not display a general growth defect in the absence of ·NO stress (Fig 2), and we confirmed that the difference in ·NO detoxification observed could be attributed to synthesis of and enzymatic detoxification by Hmp (Fig S1G & S1H). Using a GFP reporter, we demonstrated that Δ*greA*Δ*greB* populations exhibited bimodal protein expression under ·NO stress, where one subpopulation had trouble synthesizing proteins, whereas the other produced protein at levels comparable to WT (Fig 3). Bimodal protein expression was observed with multiple promoters (Fig 4), and sorting experiments confirmed that it was a transient phenotypic diversification, rather than a heritable genetic change (Fig 5). Further analyses revealed that the phenotypic diversification originated at the transcriptional level and depended on bacterial RNAP (Fig 6–8). However, we did not delineate the exact mechanism of how loss of Gre factors led to stress-induced phenotypic diversification by RNAP in this study. Given the potential of ·NO autoxidation products to damage DNA and stall RNAP, we postulate that differential damage to or repair of DNA can produce subpopulations with arrested RNAPs in the absence of Gre factors. Alternatively, given the indirect regulatory role of Gre factors on gene expression, it is possible that downstream effects of Gre factor loss prime *E. coli* for phenotypic diversification upon toxic metabolite exposure [21,22,24,27]. Given the ubiquity of Gre factors in prokaryotes and the inherently stressful lives of microbes, unraveling the molecular mechanism of how Gre factors prevent stress-induced phenotypic diversification will be a fertile area for future study. Further, previous work illustrated a role for Gre factors in limiting the epigenetic switching of the *lac* gene network under unstressed growing conditions, which suggests that Gre factors play a role in preventing diversification under a broad swath of environments [62].

Interestingly, phenotypic diversification is often hypothesized to promote bet-hedging for when pathogens encounter variable environments [63–65]. For example, enteric pathogens such as *E. coli* and *Salmonella* experience drastically different environments while invading the gastrointestinal tract [66]. The esophagus, stomach, intestines, and subepithelial tissues are unique environments with differing stress conditions, including shifts from acidic to basic conditions, high to low oxygen levels, and presence or absence of important nutrients such as iron [66]. Heterogeneity is thought to foster adaptation of populations as a whole at the cost of reduced fitness for specific subpopulations [66,67]. However, a potential caveat for the benefits of diversification is cheating in the presence of community variables. ·NO and H_2_O_2_ are highly diffusible toxic metabolites that act effectively as community poisons until their concentrations fall below inhibitory levels. Due to their high diffusivity, cells cannot propagate until the environment has been cleared of ·NO or H_2_O_2_, and thus proficient detoxifiers will commit resources to deal with more than their fair share of poison if neighbors are deficient at detoxification. Such a scenario ultimately delays the elimination of the community insult, leads to prolonged periods of stress, and negatively impacts the entire population since the most fit in those conditions do not increase in abundance compared to those that are the least fit. Indeed, those outcomes were what we observed using monocultures of Hmp-sufficient (most fit) and Hmp-deficient (least fit) cells and a coculture to model the Gre factor mutant (Fig 9). Therefore, by Gre factors protecting against phenotypic diversification under ·NO stress, they also serve to protect against cheating in such conditions.

In addition, we demonstrated that bimodal protein expression during H_2_O_2_ stress occurs in Δ*greA*Δ*greB* but not WT (Fig 10). Similar to ·NO, H_2_O_2_ is an antimicrobial produced within phagocytes that rapidly diffuses across bacterial cell membranes [41,49]. It also has broad spectrum activity and can damage similar biomolecules within bacteria such as DNA, Fe-S clusters, and thiol groups [49,68]. Microbes possess enzymatic defense systems, such as catalases and peroxidases, to combat the deleterious effects of H_2_O_2_ [41]. However, unlike bacterial ·NO defenses, H_2_O_2_ defenses are present under normal growth conditions, although their expression is also induced by H_2_O_2_ [49,69]. Under such circumstances, cheating behavior would likely manifest only under longer periods of H_2_O_2_ exposure, which would likely explain the smaller difference in clearance times that we observed between WT and Δ*greA*Δ*greB* for H_2_O_2_ (Fig. S7B) compared to ·NO (Fig. 1). Given the pervasiveness of toxic metabolites in natural and anthropogenic settings [4,70,71], the role of Gre factors in modulating cheating behavior and competition between microbes could be broadly applicable.

In conclusion, Gre factors play vital regulatory roles in transcriptional homeostasis, and their importance to the pathogenicity of numerous microbes continue to be uncovered [34–38]. The data presented here adds to the growing understanding of Gre factors and their role in bacterial stress physiology. Moreover, results from this work suggest that inhibitors of Gre factors could decrease the virulence of pathogens that rely on detoxification of toxic metabolites to foster their infectivity. Such agents would work independently of current antibiotics and since loss of Gre factors does not lead to growth inhibition under normal conditions, it would be projected that the time-scale for resistance development would be longer than conventional treatments [72,73].

## Materials and methods

### 3.4.1 Bacterial strains

All strains were derived from *E. coli* K12 MG1655 (WT). Deletion mutants were generated by P1 bacteriophage mediated transduction from the associated mutant in the Keio collection, which has been described previously, with the exception of *lacI*::*kanR*, *araBAD*::*P*_T5_-*mcherry*-*kanR* and *araBAD*::*P*_T5_-*gfp*-*kanR* that were introduced using the lambda red system [74,75]. Following P1 transduction, strains were cured of the chromosomally-inserted kanamycin resistance marker using a pCP20 plasmid, which expresses FLP recombinase [75]. For double and triple deletion mutants, consecutive rounds of P1 transduction and curing were performed. For a list of strains used in this study, refer to Table S1. All deletions were confirmed by PCR. Refer to Table S2A and S2B for a list of primers used in mutant construction and verification.

### 3.4.2 Chemicals and growth media

The growth medias used in this study were Luria Bertani (LB) broth, and MOPS minimal media and M9 minimal media, which were both supplemented with 10 mM glucose as the sole carbon source. LB broth was made by combining LB powder (40% tryptone, 20% yeast extract, 40% NaCl per gram of solid) with Milli-Q water (18.2 MΩ • cm at 25°C) and sterilizing the solution by autoclaving. MOPS and M9 minimal media were sterilized by filtration. The ·NO donor used, (Z)-1-[N-(3-aminopropyl)-N-(3-ammoniopropyl)amino]diazen-1-ium-1,2-diolate (DPTA NONOate) (Cayman Chemical), was dissolved in 10 mM NaOH to a concentration of 72 mM and stored on ice. Kanamycin, ampicillin, and isopropyl ß-D-1-thiogalactopyranoside (IPTG) were dissolved in autoclaved Milli-Q water at concentrations of 50 mg/mL, 100 mg/mL, and 2.38 g/mL, respectively, sterile-filtered and stored at 4°C prior to use. DFHBI-1T was resuspended in DMSO at a concentration of 5 mg/mL. For ·NO probe calibrations, a solution of SNAP (S-Nitroso-N-Acetyl-D,L-Penicillamine) and ethylenediaminetetraacetic acid (EDTA) was prepared by reconstituting 5.6 mg SNAP and 5 mg EDTA in 25 mL autoclaved Milli-Q water. A 0.1 M CuCl_2_ solution was prepared by dissolving 8.5 g CuCl_2_·2H_2_O in 500 mL of Milli-Q water. For H_2_O_2_ treatment assays, an H_2_O_2_ solution (35% wt solution in H_2_O – Fisher Scientific) was dissolved in autoclaved Milli-Q water at a concentration of 25, 50, or 75 mM.

### 3.4.3 Plasmids

For a list of plasmids used in this study refer to Table S1. All plasmids were derived from either a low-copy (pUA66) or high-copy plasmid (pQE80), and the parental plasmid for each construct is specified in Table S1. pJR05, pSA21, pXW02 and pXW09 were obtained from previous works. pDS01 and pDS02 were constructed by restriction digestion with BamHI and SbfI followed by ligation with a Quick Ligation Kit (NEB). All other plasmids were constructed using a HiFi DNA assembly kit (NEB). All plasmids were confirmed by Sanger sequencing (Genewiz). Refer to Table S2C and S2D for a list of primers and double stranded DNA sequences used plasmid construction.

### 3.4.4 ·NO measurements

·NO concentrations in bioreactors were continuously measured using an ·NO-sensing electrode (World Precision Instruments). Daily calibrations were performed by submerging the probe in a solution of 10 mM CuCl_2_ and adding increasing doses of SNAP. Raw signal generated from the probe, in units of picoamps, were converted to units of ·NO concentration (in μM) using a scaling factor of 0.457 ·NO per molecule of SNAP, which was determined previously [18].

### 3.4.5 H_2_O_2_ measurements

H_2_O_2_ concentrations were measured using Amplex Red hydrogen peroxide/peroxidase kits (Life Technologies) according to the manufacturer’s instructions. A standard curve with concentrations of 0, 1, 2.5, 5 and 10 μM H_2_O_2_ was constructed. Samples were diluted to a concentration within 10 μM to convert fluorescence values to H_2_O_2_ concentrations.

### 3.4.6 Absorbance and fluorescence measurements

Cell density was measured during experiments by sampling 300 μL of cultures from bioreactors or flasks, and measuring the absorbance at 600 nm using a 96-well clear, flat bottom plate (Corning) and a Synergy^™^ H1 Hybrid Microplate Reader. Fluorescence measurements were performed on a LSRII flow cytometer (BD Bioscience). For GFP and tdBroccoli, an excitation wavelength of 488 nm and emission bandpass filter of 525/550 was used. For mCherry, the excitation wavelength and emission bandpass filter were set to 561 and 610/620 respectively. For each sample, 50,000 cellular events were recorded.

### 3.4.7 ·NO and H_2_O_2_ consumption assays

*E. coli* were taken from a −80°C frozen stock, inoculated into 1 mL of LB media and grown for 6 hr in an incubator at 37°C and 250 RPM. After 6 hr, 10 μL of culture were transferred to 1 mL of MOPS media and incubated for 16 hr at 37°C and 250 RPM. After 16 hr, OD_600_ were measured and corresponding volumes of culture were transferred into 20 mL MOPS media in 250 mL baffled flasks to achieve OD_600_ of 0.01. Flasks were incubated at 37°C and 250 RPM until cultures reached mid-exponential phase (OD_600_ = 0.2). Then, 8 mL of culture were transferred to 8 prewarmed (37°C) microcentrifuge tubes, and spun at 15,000 RPM for 3 min. Nine hundred-eighty μL of supernatant were removed from each tube and cell pellets were resuspended in 1 mL of pre-warmed MOPS media.

For ·NO consumption assays, concentrated cultures were used to inoculate bioreactors containing 10 mL MOPS media to an OD_600_ of 0.05. Immediately after inoculation, DPTA NONOate was added to an initial concentration of 250 μM. For assays involving Hmp-GFP translational fusions, the concentration of DPTA was increased to 350 μM to achieve similar ·NO concentration profiles as WT cultures. Throughout assays, ·NO concentrations were continuously monitored. Cell density measurements were taken by extracting 300 μL from bioreactors at indicated time points and measuring the OD_600_.

For H_2_O_2_ consumption assays, M9 growth media was used instead of MOPS minimal media, as MOPS had previously been shown to interfere with Amplex Red H_2_O_2_ quantification [76]. Immediately before inoculation, H_2_O_2_ was added to reactors to the desired concentrations (25, 50, or 75 μM). Afterward, concentrated cultures were used to inoculate bioreactors to an OD_600_ of 0.01. At each time point, 300 μL of culture were removed and sterile filtered using a 0.22 μM syringe filter (Millex). Samples for initial time points (t=0) were removed prior to inoculation with cells. All samples were stored on ice prior to quantification of [H_2_O_2_].

In experiments involving strains harboring plasmids, the appropriate concentrations of antibiotic (50 μg/mL kanamycin or 100 μg/mL ampicillin) were included in growth media at every step.

### 3.4.8 Cell survival assay

Cells were prepared following the same procedure as ·NO and H_2_O_2_ consumption assays. After inoculating cells into the bioreactor, 300 μL samples were removed to measure the initial OD_600_ and 10 μL of the 300 μL sample were serially diluted in PBS and plated onto LB-agar. Immediately after, DPTA was added to the bioreactor at a concentration of 250 μM. For subsequent time points, the same procedure was used to plate cells. Plates were incubated at 37°C for 16 h after which the number of colonies were counted and converted to CFU/mL.

### 3.4.9 Fluorescent reporter assays

Cells were prepared following the same procedure as ·NO and H_2_O_2_ consumption assays. For assays involving H_2_O_2_, 25, 50, or 75 μM of H_2_O_2_ was added prior to cells. After inoculating cells into the bioreactor, 300 μL samples were removed to measure the initial OD_600_ and a second 300 μL sample was removed and centrifuged at 15,000 RPM for 3 min. After centrifugation, 250 μL of supernatant were removed and pellets were resuspended in 250 μL of 4% paraformaldehyde in PBS (4% PFA). Samples were kept on ice for 30 min, after which they were centrifuged for 3 min at 15,000 RPM, 250 μL of supernatant were removed, the pellet was resuspended in 550 μL PBS, and then stored at 4°C until flow cytometry. After removal of initial samples (t = 0), for ·NO stress assays, 250 μM DPTA was immediately added to the bioreactor. For cells harboring IPTG-inducible plasmids, 1 mM IPTG was also added to the bioreactor at that time. For subsequent time points, 300 μL were extracted from bioreactors and the same procedure was used to prepare samples for flow cytometry. Samples were analyzed on a LSRII flow cytometer (BD Biosciences).

RNA aptamer-based assays followed the same procedure with a few modifications. DFHB-1T was added to the bioreactor at t = 0 to a final concentration of 50 μM. Samples were not fixed in 4% PFA, but instead centrifuged at 15,000 RPM for 1 min, 250 μL of supernatant was removed, pellets were resuspended in ice-cold PBS containing 50 μM DFHB-1T, and stored on ice prior to flow cytometry. Samples were analyzed on the same day, immediately after collection of the terminal timepoint. For mCherry-tdBroccoli (Fig 7), 2 mM IPTG was introduced (instead of 1 mM) and terminal samples were removed at t = 5 hr (instead of 3 hr), to increase fluorescence output and allow the signal to further develop.

For Figures 2B and S1F, experiments were performed in 250 mL baffled flasks as opposed to bioreactors. Cultures were inoculated at an OD_600_ of 0.01, and after 2 hr of incubation at 37°C and 250 RPM, 300 μL samples were removed and prepared for flow cytometry following the procedure outlined above. Immediately after, 1 mM IPTG was added to flasks. Samples were removed every hr for 3 hr and prepared for flow cytometry.

### 3.4.10 Cell sorting assay

Cells were prepared following the same procedure as ·NO and H_2_O_2_ consumption assays. At t = 0, 250 μM DPTA and 1 mM IPTG were immediately added to the bioreactor. After 1 hr, 1 mL samples were removed, spun at 15,000 RPM for 3 min, 950 μL of supernatant were removed, and pellets were resuspended in 1950 μL ice-cold PBS and stored on ice. Sample were sorted on a FACS ARIA fusion (BS Biosciences) in which 1 million high-GFP cells (top 10^th^ percentile), 1 million low-GFP cells (bottom 10^th^ percentile), and 1 million cells from the total distribution were collected in microcentrifuge tubes and stored on ice. Collected samples were centrifuged at 15,000 RPM for 3 min, 950 μL of supernatant were removed, and pellets were resuspended in 950 μL of LB containing 50 μg/mL kanamycin. Samples were transferred to test tubes and incubated at 37°C and 250 RPM for 12 hr. Afterwards, 500 μL of sample was mixed with 500 μL of 50% glycerol in cryotubes and stored at −80°C. For culturability measurements after cell sorting (Fig S4A & S4B), 500,000 cells were collected instead, and sorted samples were immediately diluted in PBS and plated onto LB-agar containing 50 μg/mL kanamycin.

### 3.4.11 Co-culture experiments

Cultures of Hmp-proficient (mCherry) and Hmp-deficient (GFP) were prepared concurrently using the same procedure as ·NO and H_2_O_2_ consumption assays. Both strains were inoculated in a 1:1 ratio in a bioreactor at an OD_600_ of 0.025. A 300 μL sample was removed to measure the initial OD_600_ and the same sample was centrifuged at 15,000 RPM for 3 min. After centrifugation, 250 μL of supernatant were removed and pellets were resuspended in 250 μL of 4% paraformaldehyde in PBS (4% PFA). Samples were kept on ice for 30 min, after which they were centrifuged for 3 min at 15,000 RPM, 250 μL of supernatant were removed, pellets were resuspended in 550 μL PBS, and then stored at 4°C until flow cytometry. After removal of initial samples (t = 0), 250 μM DPTA was immediately added to the bioreactor and the ·NO concentration of the reactor was continuously monitored. For subsequent time points, 300 μL were extracted for OD_600_ measurements and the same procedure was used to prepare samples for flow cytometry. Samples were analyzed on a LSRII flow cytometer (BD Biosciences) and the proportion of mCherry positive and GFP-positive cells were quantified. The OD_600_ of the Hmp-proficient subpopulation and Hmp-deficient subpopulation were calculated by multiplying the total OD_600_ by the respective fraction of fluorescent cells. OD_600_ values were normalized by dividing all values by the initial OD_600_ for each subpopulation. Initial co-culture experiments were performed at an OD_600_ of 0.05 (Fig S8); however, differences in ·NO detoxification between Hmp-proficient monocultures and cocultures were not large enough to detect different in growth resumption due to the different time resolutions of those measurements. To address this hurdle, we lowered the initial OD_600_ to 0.025, to better observe differences in ·NO detoxification and growth.

## Acknowledgments

We would like to thank Wen Kang Chou, Heather Cho, Christina J. DeCoste, and Katherine Rittenbach for assistance, as well as the National BioResource Project (NIG, Japan) for distribution of the Keio collection.

