## Supplemental Figure 1 for "Gre factors protect against phenotypic diversification and cheating in *Escherichia coli* populations under toxic metabolite stress"

### Figure S1

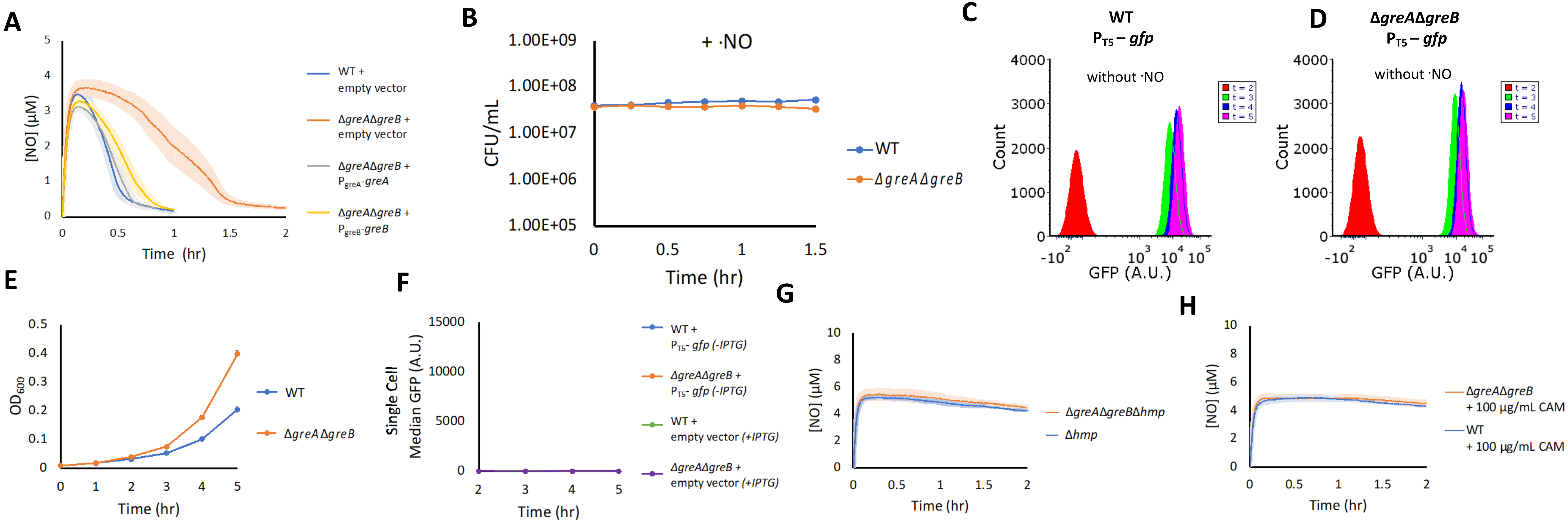


**(A) Complementation of *greA* or *greB* restores ·NO detoxification to WT levels.** Cultures were grown in MOPS minimal media to mid-exponential phase and inoculated into a bioreactor at an $OD_{600}$ of 0.05. Immediately after, 250 µM of DPTA NONOate was added to the bioreactor. ·NO concentrations were continuously monitored in the bioreactor. WT harboring an empty puA66 plasmid (blue), Δ*greA*Δ*greB* harboring an empty puA66 plasmid (orange), Δ*greA*Δ*greB* harboring puA66 expressing *greA* under its native promoter (grey). Δ*greA*Δ*greB* harboring puA66 expressing *greB* under its native promoter (yellow). Solid lines represent the means of at least three independent replicates, whereas light shading represents the standard errors of the means.

**(B) Culturability under ·NO stress.** Cultures were grown in MOPS minimal media to mid-exponential phase and inoculated into a bioreactor at an $OD_{600}$ of 0.05. Immediately after, 250 µM of DPTA NONOate was added to the bioreactor. Samples were removed at the indicated time points, diluted in PBS, plated on LB agar plates, and incubated at 37°C. After 16 hours, colonies were counted. WT (blue), Δ*greA*Δ*greB* (orange). Circles represent the means of at least three replicates, whereas error bars represent the standard errors of the means. For both WT and Δ*greA*Δ*greB*, CFU measurements were not significantly different than the respective CFU measurements at t = 0, using a t-test and a significance threshold of $p < 0.05$.

**(C) & (D) GFP distributions in the absence of ·NO stress,** WT and Δ*greA*Δ*greB* cultures were inoculated into baffled flasks containing MOPS minimal media at and $OD_{600}$ of 0.01 and incubated at 37 °C and 250 RPM. After 2 hrs of incubation, 1 mM IPTG was added to flasks to induce GFP production. Samples were removed, fixed, and GFP was measured by flow cytometry. Images are representative of at least 3 biological replicates.

**(E) & (F) Δ*greA*Δ*greB* display a growth advantage independent of the $P_{T5}$-gfp vector and GFP is negligible in the absence of either $P_{T5}$-gfp vector or IPTG.** (E) WT (blue) and Δ*greA*Δ*greB* (orange) cultures were inoculated into baffled flasks containing MOPS minimal media at and $OD_{600}$ of 0.01 and incubated at 37 °C and 250 RPM. Samples were removed to measure $OD_{600}$ at indicated time points. (F) Cultures were inoculated into baffled flasks containing MOPS minimal media at and $OD_{600}$ of 0.01 and incubated at 37 °C and 250 RPM. After 2 hrs of incubation, either 1 mM IPTG was added (+IPTG) or an equivalent volume of sterile DI water (-IPTG) was added to flasks. Samples were removed at indicated time points, fixed, and GFP was measured by flow cytometry. WT $P_{T5}$-*gfp* without IPTG (blue), Δ*greA*Δ*greB* $P_{T5}$-*gfp* without IPTG (orange), WT + empty vector with IPTG (green), Δ*greA*Δ*greB* + empty vector with IPTG (purple). Data is plotted with the same axis as Figure 2B. Circles represent the means of at least 3 independent replicates. Error bars represent the standard errors of the means.

**(G) & (H) Difference between Δ*greA*Δ*greB* and WT·NO detoxification requires *hmp* and de novo protein synthesis.** Cultures were grown in MOPS minimal media to mid-exponential phase and inoculated into a bioreactor at an $OD_{600}$ of 0.05. Immediately after, 250 µM of DPTA NONOate was added to the bioreactor and ·NO concentrations were continuously monitored. (G) Δ*hmp* (blue) and Δ*greA*Δ*greB*Δ*hmp* (orange). (H) WT (blue) and Δ*greA*Δ*greB* (orange) cultures were treated with 100 µg/mL chloramphenicol (CAM) for 15 min prior to and during DPTA treatment. Solid lines represent the means of at least three independent replicates, whereas light shading represents the standard errors of the means.
