## Supplemental Figure 2 for "Gre factors protect against phenotypic diversification and cheating in *Escherichia coli* populations under toxic metabolite stress"

Figure S2

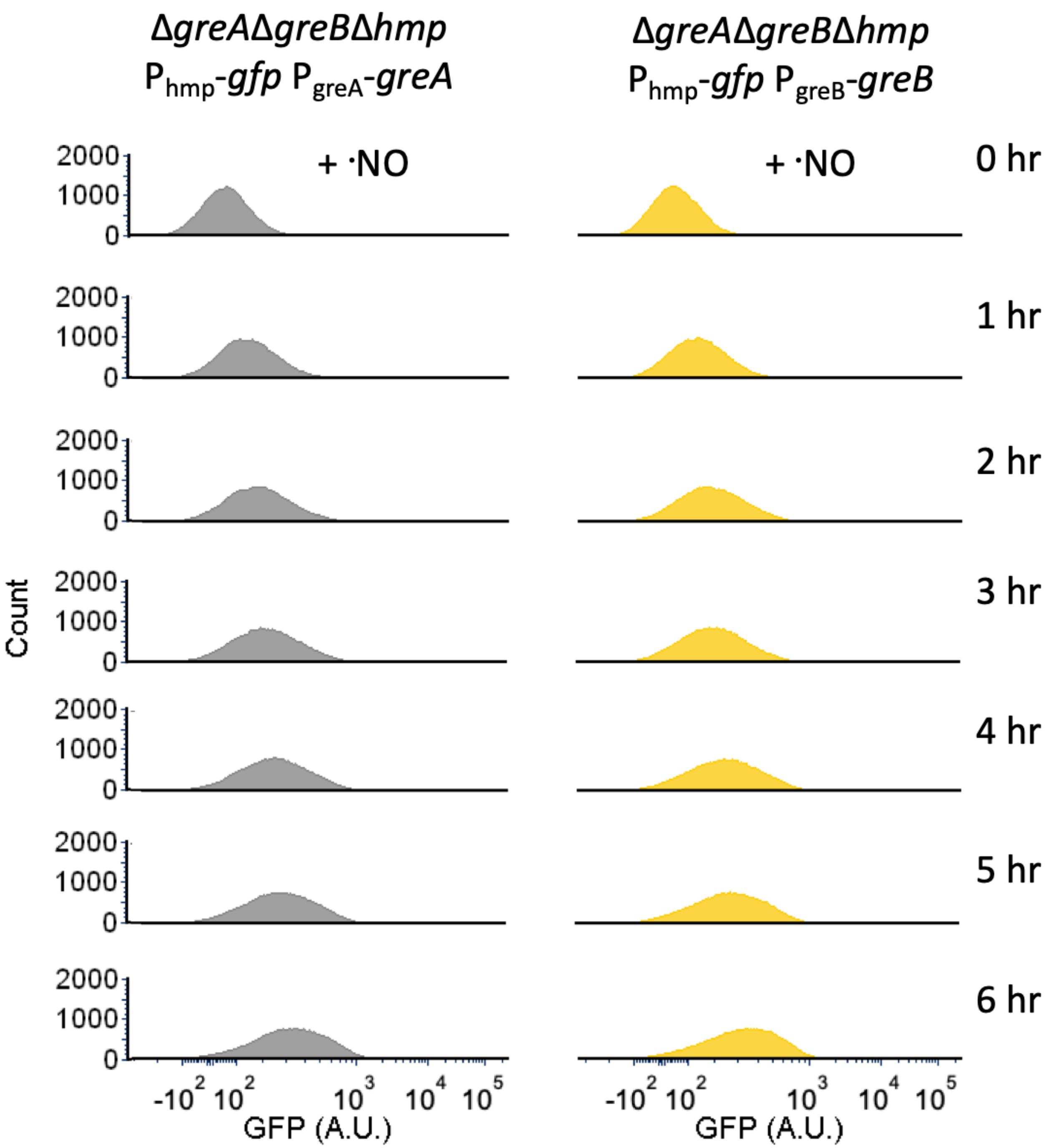


**Figure S2. Complementation of *greA* or *greB* restores unimodal fluorescence from the *hmp* promoter**

Cultures were grown in MOPS minimal media to mid-exponential phase and inoculated into a bioreactor at an $OD_{600}$ of 0.05. Immediately after, 250 µM of DPTA NONOate and 1 mM IPTG were added. Samples were removed, fixed, and GFP distributions were measured by flow cytometry at the indicated time points. Fluorescence measurement were taken for additional time points to allow signal to develop. Δ*greA*Δ*greB*Δ*hmp* harboring pUA66 expressing *gfp* from the *hmp* promoter and *greA* under its native promoter (grey). Δ*greA*Δ*greB*Δ*hmp* harboring pUA66 expressing *gfp* from the *hmp* promoter and *greB* under its native promoter (yellow). Images are representative of at least 3 biological replicates.
