## Supplemental Figure 3 for "Gre factors protect against phenotypic diversification and cheating in *Escherichia coli* populations under toxic metabolite stress"

Figure S3

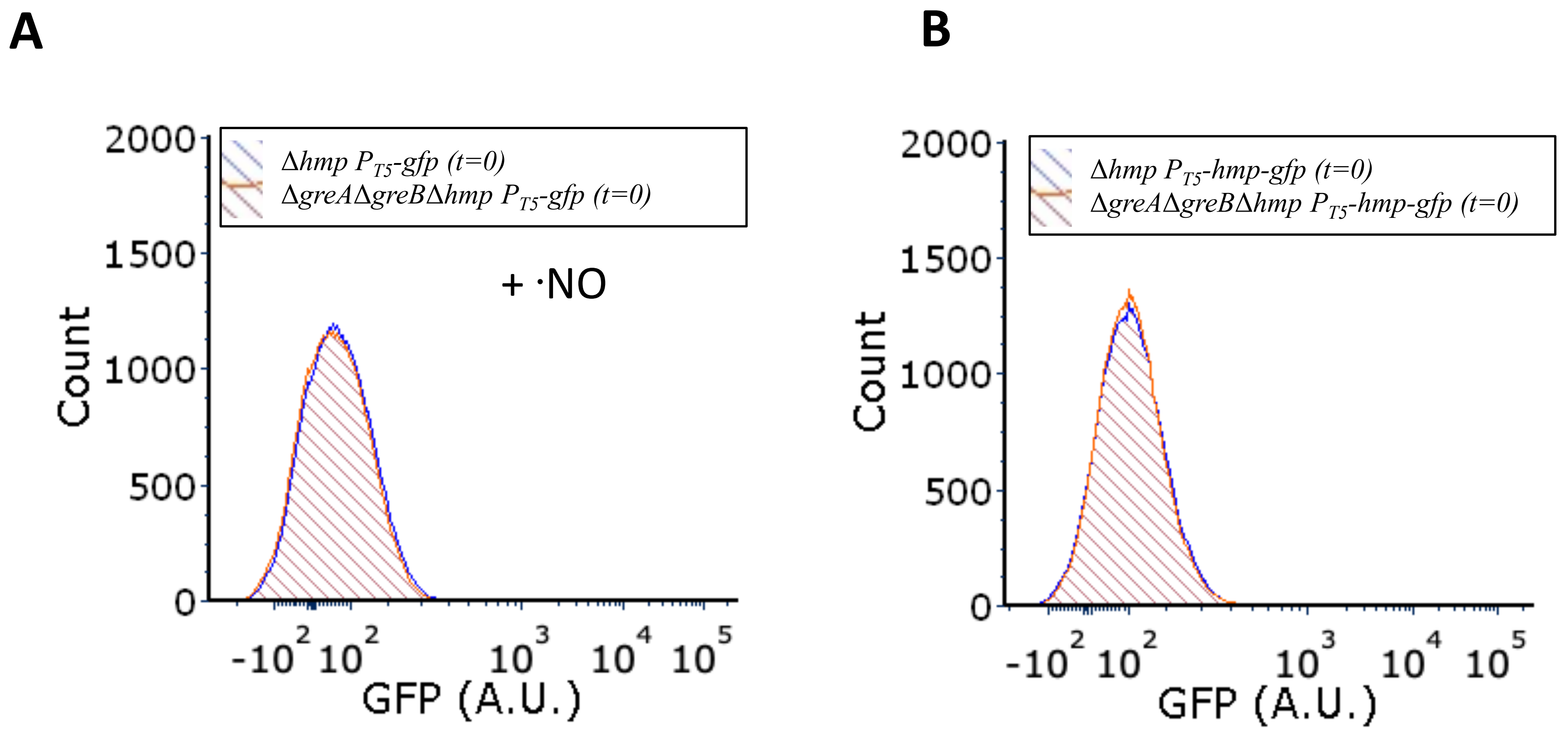


**Figure S3. Fluorescence distribution profiles prior to treatment from the $T_5$ promoter**

Cultures were grown in MOPS minimal media to mid-exponential phase and inoculated into bioreactors at an $OD_{600}$ of 0.05. Prior to treatment (Fig 4), samples were removed (t=0), fixed, and fluorescence was measured by flow cytometry. Panel (A) reflects baseline fluorescence distributions for cells expressing GFP under the $T_5$ promoter. Panels (B) reflect baseline fluorescence distributions for cells expressing Hmp-GFP translational fusions under the $T_5$ promoter. Δ*hmp* (blue) and Δ*greA*Δ*greB*Δ*hmp* (orange). Fluorescent histograms are representative of three or more biological replicates.
