## Supplemental Figure 4 for "Gre factors protect against phenotypic diversification and cheating in *Escherichia coli* populations under toxic metabolite stress"

### Figure S4

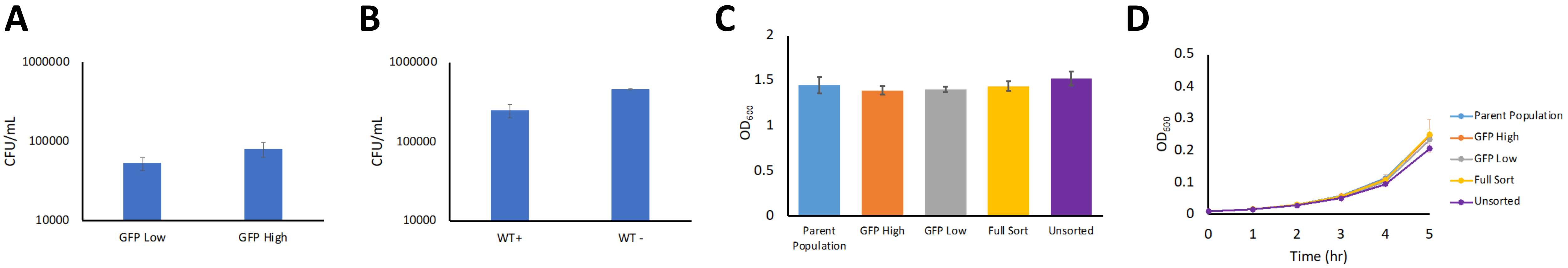


**(A) & (B) Culturability of Δ*greA*Δ*greB*Δ*hmp* and *Δhmp* GFP populations after cell sorting.** (A) Δ*greA*Δ*greB*Δ*hmp* (parent population) harboring a $P_{T5}$-*gfp* expression vector was grown in MOPS minimal media to mid-exponential phase and inoculated into bioreactors at an $OD_{600}$ of 0.05. Immediately after, 250 µM of DPTA NONOate and 1 mM IPTG were added to the bioreactor. A sample was removed at 1 hr and passed through a cell sorter to collect 500,000 cells from low and high GFP peaks. Collected cells were diluted in PBS and plated on LB agar containing 50 µg/mL kanamycin for plasmid retention. Data represent the means of three independent replicates, whereas the error bars represent the standard errors of the means. We note that culturability following sorting for Δ*greA*Δ*greB*Δ*hmp* in these conditions was low, and we assayed to what extent ·NO contributes to that observation. (B) Δ*hmp* harboring a $P_{T5}$-*gfp* expression vector was grown in MOPS minimal media to mid-exponential phase and inoculated into bioreactors at an $OD_{600}$ of 0.05. Immediately after, either 1 mM IPTG (WT-) or 250 µM of DPTA NONOate and 1 mM IPTG (WT+) were added to the bioreactor. Samples were removed at 1 hr and passed through a cell sorter to collect 500,000 GFP positive cells. Collected cells were diluted in PBS and plated on LB agar containing 50 µg/mL kanamycin for plasmid retention. Data represent the means of two independent replicates, whereas error bars represent the standard errors of the means. These data suggest that ·NO reduces the culturability of cells after sorting, and that Δ*greA*Δ*greB*Δ*hmp* has lower culturability than Δ*hmp* in these sorting conditions.

**(C) & (D) Δ*greA*Δ*greB*Δ*hmp* subpopulations have comparable terminal cell densities and growth dynamics.** Δ*greA*Δ*greB*Δ*hmp* (parent population) harboring a $P_{T5}$-*gfp* expression vector was grown in MOPS minimal media to mid-exponential phase and inoculated into bioreactors at an $OD_{600}$ of 0.05. Immediately after, 250 µM of DPTA NONOate and 1 mM IPTG were added to the bioreactor. A sample was removed at 1 hr and passed through a cell sorter to collect 1 million cells from the bottom 10 percent (GFP low), top 10 percent of the fluorescence distribution (GFP high), and total distribution (Full sort). One million cells were also collected from the original sample, without passing through the cell sorter (Unsorted). Collected cells were recovered in rich media, mixed in a 1:1 ratio with 50% glycerol, and stored at -80°C. Those -80°C frozen stocks were then inoculated in 1 mL LB media and grown for 6 hours in an incubator at 37°C and 250 RPM. Ten µL were then transferred into 1 mL MOPS minimal media and grown for 16 hrs at 37°C and 250 RPM. Panel (C): the terminal cell density of cultures after 16 hrs of incubation. Panel (D): A corresponding volume of culture was transferred into 20 mL MOPS media in a 250 mL baffled flask to reach an $OD_{600}$ of 0.01. Flasks were incubated at 37°C and 250 RPM and $OD_{600}$ was measured every hr for 5 hours. Bars and colored circles represent the means of at least three independent replicates, whereas error bars represent the standard errors of the means.
