## Supplemental Figure 5 for "Gre factors protect against phenotypic diversification and cheating in *Escherichia coli* populations under toxic metabolite stress"

Figure S5

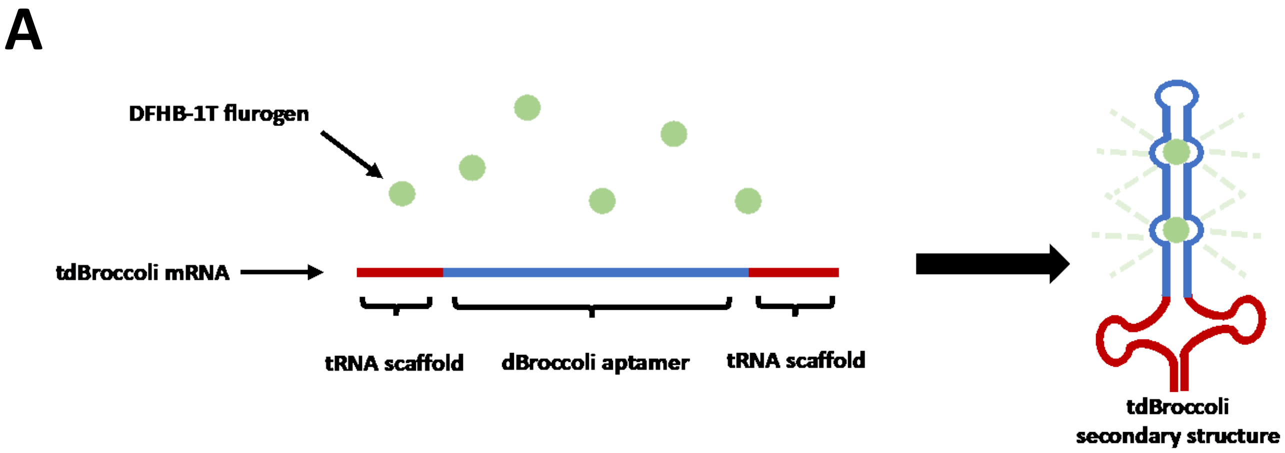


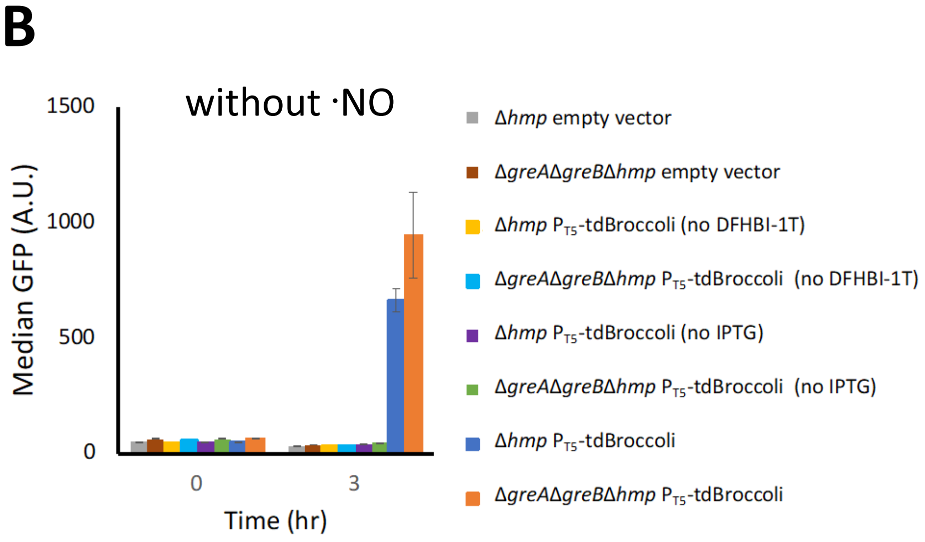


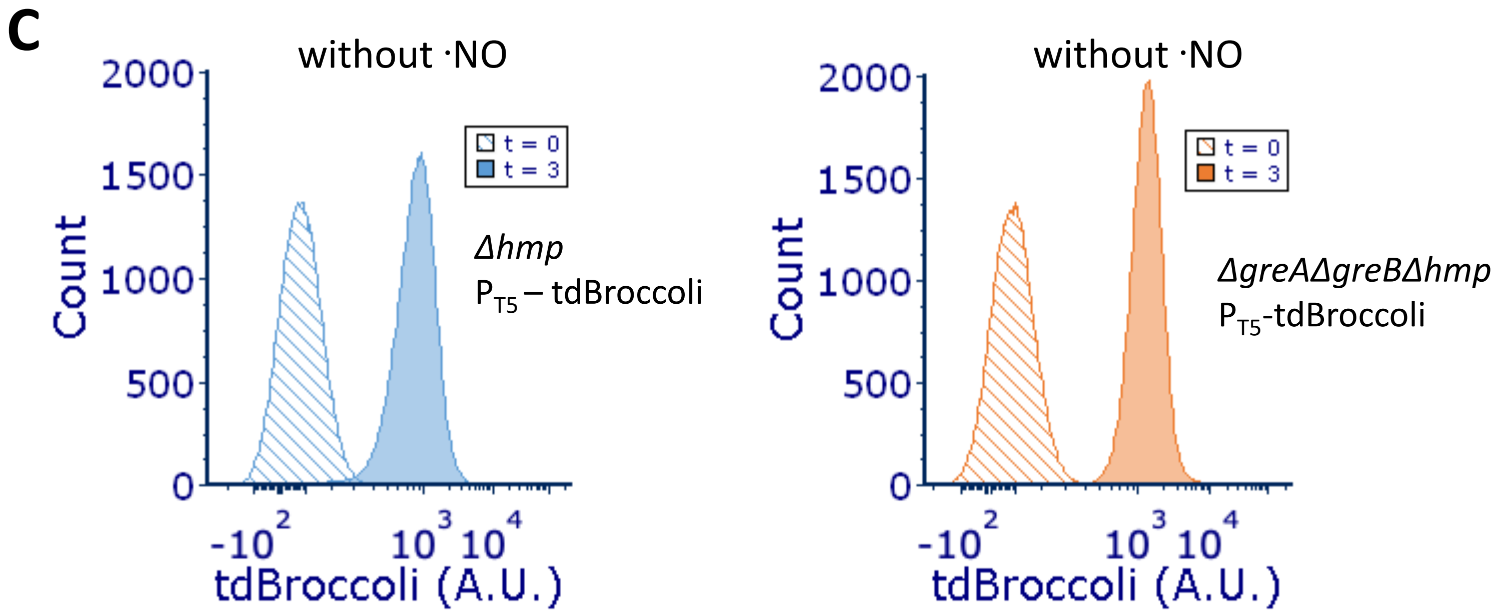


**Figure S5. tdBroccoli fluorescence measurements in the absence of ·NO**

(A) A cartoon depiction of the tdBroccoli RNA aptamer expression system. Upon transcript synthesis, a secondary structure is formed that accommodates the fluorogen DFHBI-T. Cultures were grown in MOPS minimal media to mid-exponential phase and inoculated into a bioreactor at an $OD_{600}$ of 0.05. Immediately after, 1 mM IPTG, and 50 µM DFHBI-1T were added to the bioreactor, unless otherwise stated. Samples were removed at indicated time points, and fluorescence was measured by flow cytometry. (B) Median GFP fluorescence measurements. Solid bars represent the means of at least three independent replicates, whereas error bars represent the standard errors of the means. (C) Fluorescence distributions. Images are representative of at least three biological replicates.
