## Supplemental Figure 6 for "Gre factors protect against phenotypic diversification and cheating in *Escherichia coli* populations under toxic metabolite stress"

Figure S6

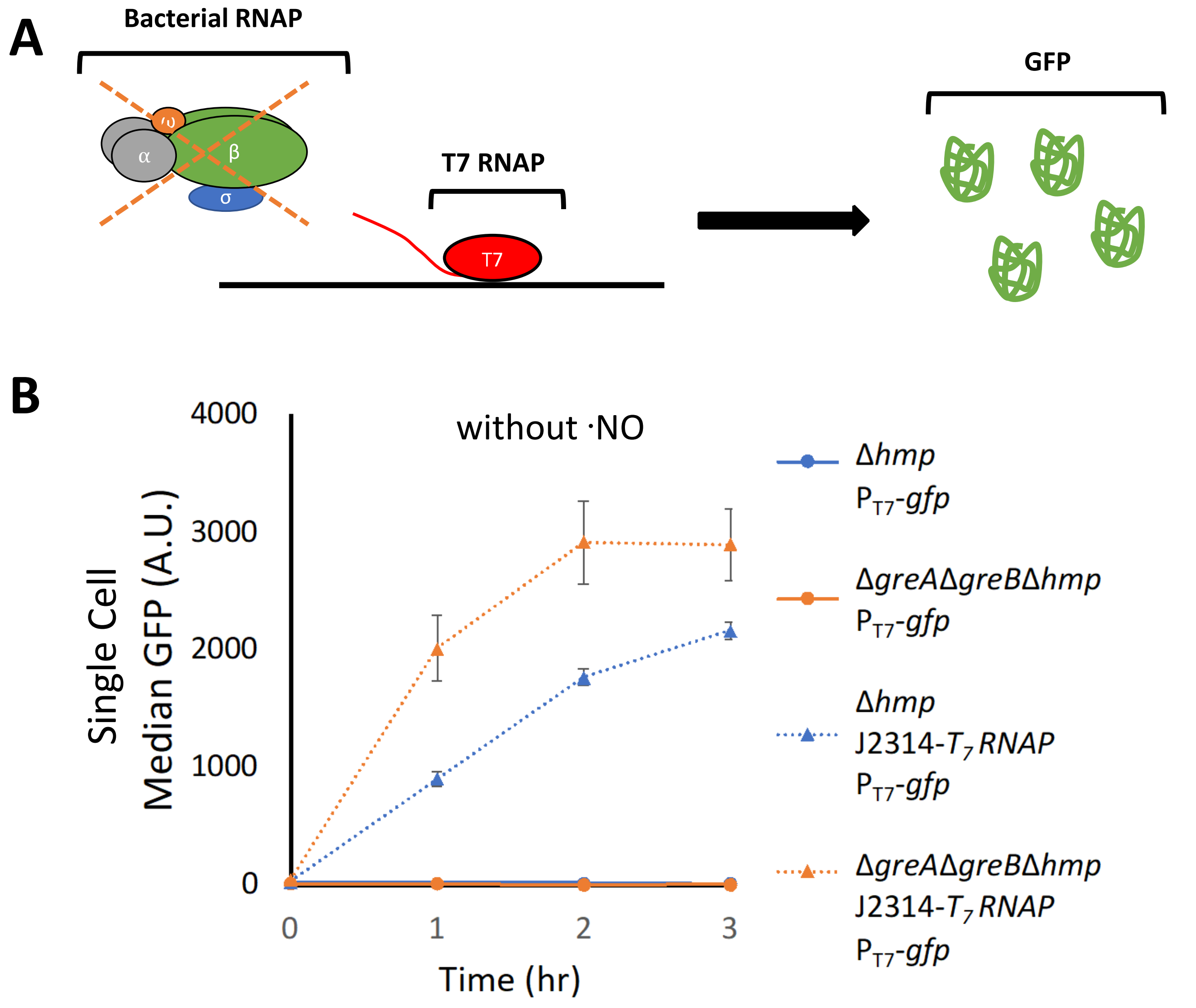


**Figure S6. GFP expression from the $T_7$ promoter requires $T_7$ RNA polymerase**

(A) Cartoon depiction of the $T_7$ expression system constructed. Bacterial RNAP was inhibited through the introduction of rifampicin, which allows GFP expression to be solely under the control of previously translated, constitutively expressed $T_7$ RNAP. (B) Cultures were grown in MOPS minimal media to mid-exponential phase and inoculated into a bioreactor at an $OD_{600}$ of 0.05. Immediately after, 1 mM IPTG was added to the bioreactor. Samples were removed at indicated time points, fixed and median GFP fluorescence was measured by flow cytometry. Data represent the means of three independent replicates with error bars representing the standard errors of the means. Δ*hmp* (blue) and Δ*greA*Δ*greB*Δ*hmp* (orange). Cells harboring a plasmid containing $P_{T7}$-*gfp* are depicted as circles with solid lines. Cells harboring a plasmid containing $P_{T7}$-*gfp* and J2314-*$T_7$ RNAP* are depicted as triangles with dashed lines.
