## Supplemental Figure 7 for "Gre factors protect against phenotypic diversification and cheating in *Escherichia coli* populations under toxic metabolite stress"

Figure S7

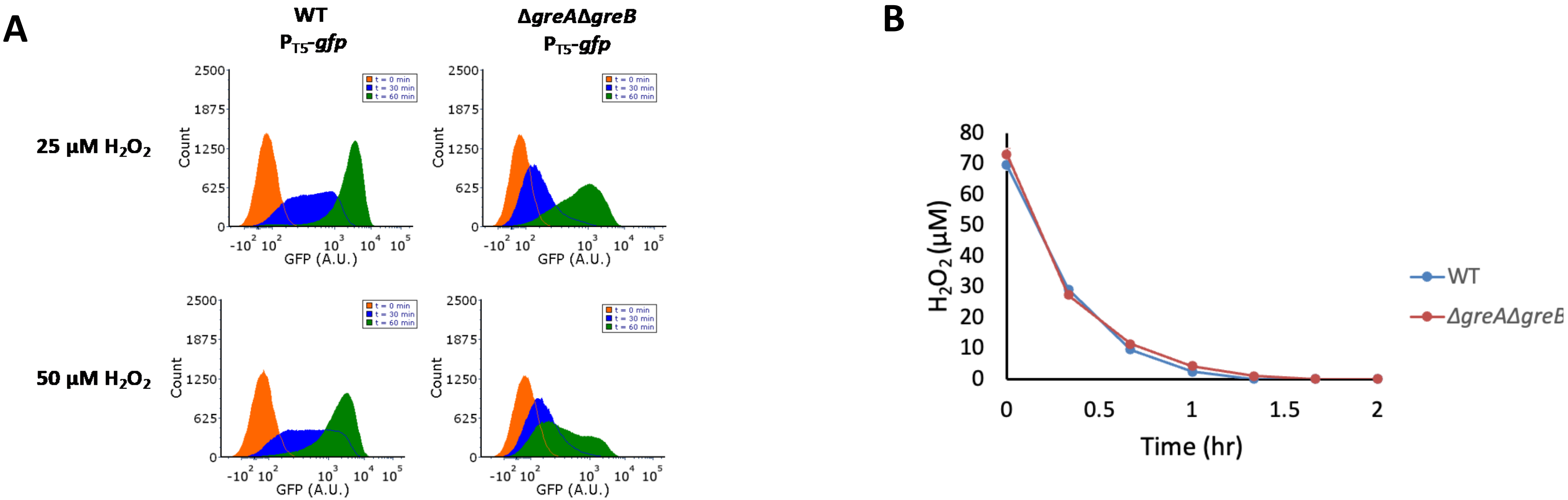


**(A) GFP distributions under 25 µM and 50 µM $H_2O_2$ treatment.** Cultures were grown in M9 minimal media to mid-exponential phase and inoculated into a bioreactor containing either 25 or 50 µM $H_2O_2$. Immediately after 1 mM IPTG was added. Samples were removed at 0, 30 and 60 min, fixed and GFP distributions were measured by flow cytometry. Images are representative of at least three biological replicates.

**(B) $H_2O_2$ clearance after 75 µM $H_2O_2$ treatment.** Cultures were grown in M9 minimal media to mid-exponential phase and inoculated into a bioreactor containing 75 µM $H_2O_2$. $H_2O_2$ concentrations were quantified every 20 min for 2 hrs. $H_2O_2$ fell to 0 µM at 1.33 hr (80 min) for WT, and 1.66 hr (100 min) for Δ*greA*Δ*greB*. Solid circles represent the means of three independent replicates, whereas error bars represent the standard errors of the means. WT (blue) and Δ*greA*Δ*greB* (red).
