## Supplemental Figure 8 for "Gre factors protect against phenotypic diversification and cheating in *Escherichia coli* populations under toxic metabolite stress"

Figure S8

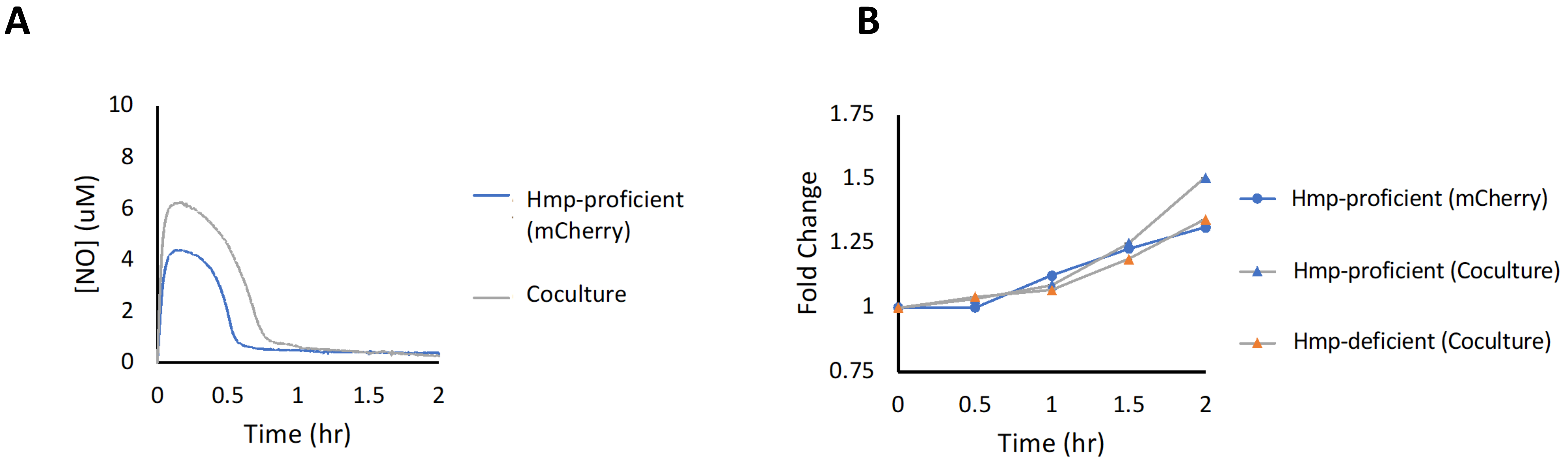


**Figure S8. ·NO consumption and outgrowth at an $OD_{600}$ of 0.05**

Cultures were grown in MOPS minimal media to mid-exponential phase and inoculated into a bioreactor at an $OD_{600}$ of 0.05 either as a monoculture of Hmp-proficient cells (blue) or as a 1:1 coculture of Hmp-proficient cells and Hmp-deficient cells (grey). Immediately after, 250 µM of DPTA NONOate was added to the bioreactor.

(A) ·NO concentrations were continuously monitored in the bioreactor. Solid lines represent a single replicate.

(B) Samples were removed to measure $OD_{600}$ at indicated time points. For co-cultures, samples were fixed and the $OD_{600}$ was scaled by the proportion of GFP and mCherry fluorescent cells estimated by flow cytometry. Colored circles and triangles represent a single replicate.
