## Supplemental Table 1 for "Gre factors protect against phenotypic diversification and cheating in *Escherichia coli* populations under toxic metabolite stress"

### Table S1. List of bacterial strains and plasmids

| Strain | Genotype | Notes |
|---|---|---|
| **MG1655** | $F^-$ $\lambda^-$ *ilvG*$^-$ *rfb-50 rph-1* | ATCC 700926 [1] |
| **MO001** | MG1655 Δ*lacI::lacIq* Δ*lacZYA::*$P_{T5}$- *mcherry-kanR* | [2] |
| **MO002** | MG1655 Δ*lacI::lacIq* Δ*lacZYA::*$P_{T5}$- *mcherry* | [3] |
| **WM020** | MG1655 Δ*lacI::lacIq* Δ*lacZYA::*$P_{T5}$- *gfp-kanR* | [4] |
| **DS001** | MG1655 Δ*hmp::kanR* | P1 transduction of mutation in Keio collection into MG1655 |
| **DS002** | MG1655 Δ*greA::kanR* | P1 transduction of mutation in Keio collection into MG1655 |
| **DS003** | MG1655 Δ*greB::kanR* | P1 transduction of mutation in Keio collection into MG1655 |
| **DS004** | MG1655 Δ*hmp* | *kanR* cured from DS001 |
| **DS005** | MG1655 Δ*greA* | *kanR* cured from DS002 |
| **DS006** | MG1655 Δ*greB* | *kanR* cured from DS003 |
| **DS007** | MG1655 Δ*greB*Δ*greA::kanR* | P1 transduction of mutation in Keio collection into DS003 |
| **DS008** | MG1655 Δ*greB*Δ*greA* | *kanR* cured from DS007 |
| **DS009** | MG1655 Δ*greB*Δ*greA*Δ*hmp::kanR* | P1 transduction of mutation in Keio collection into DS008 |
| **DS010** | MG1655 Δ*greB*Δ*greA*Δ*hmp* | *kanR* cured from DS009 |
| **DS011** | MG1655 Δ*lacI::kanR* | *lacI* deleted with lambda red system |
| **DS012** | MG1655 Δ*lacI* | *kanR* cured from DS011 |
| **DS013** | MG1655 Δ*lacI* Δ*hmp::kanR* | P1 transduction of mutation in Keio collection into DS012 |
| **DS014** | MG1655 Δ*lacI*Δ*hmp* | *kanR* cured from DS013 |
| **DS013** | MG1655 Δ*lacI*Δ*araBAD*::$P_{T5}$-*mcherry-kanR* | $P_{T5}$-*mcherry-kanR* knocked into DS012 with lambda red system. Used as Hmp-proficient strain in coculture. |
| **DS014** | MG1655 Δ*lacI*Δ*hmp*Δ*araBAD*::$P_{T5}$-*gfp-kanR* | $P_{T5}$-*gfp-kanR* knocked into DS014 with lambda red system. Used as Hmp-deficient strain in coculture. |

| Plasmid | Genotype | Notes |
|---|---|---|
| **pUA66** | Vector, SC101ori, *kanR*, *gfpmut2* reporter | [5] |
| **pJR05** | pUA66 $P_{T5}$-*hmp*-$gfp_{sf}$, *lacIq* | [1] |
| **pSA21** | pUA66 $P_{T5}$-$gfp_{sf}$, *lacIq* | [6] |
| **pXW02** | pUA66 $P_{hmp}$-$gfp_{sf}$ | [7] |
| **pXW09** | pQE80 $P_{T5}$-$gfp_{sf}$, *lacIq* | [7] |
| **pDS01** | pUA66 $P_{greA}$-*greA* | BamHI SbfI restriction digest and ligation |
| **pDS02** | pUA66 $P_{greB}$-*greB* | BamHI SbfI restriction digest and ligation |
| **pDS03** | pUA66 $P_{T7}$-*gfp*, *lacIq* | Hifi DNA assembly |
| **pDS04** | pUA66 J23114 – $T_7RNAP$, $P_{T7}$-*gfp*, *lacIq* | Hifi DNA assembly |
| **pDS05** | pQE80 $P_{T5}$- tdBroccoli, *lacIq* | Hifi DNA assembly |
| **pDS06** | pQE80 $P_{T5}$- *mcherry*- tdBroccoli , *lacIq* | Hifi DNA assembly |
| **pDS07** | pUA66 $P_{hmp}$-$gfp_{sf}$, $P_{greA}$-*greA* | Hifi DNA assembly |
| **pDS08** | pUA66 $P_{hmp}$-$gfp_{sf}$, $P_{greB}$-*greB* | Hifi DNA assembly |
