## Supplemental Table 2 for "Gre factors protect against phenotypic diversification and cheating in *Escherichia coli* populations under toxic metabolite stress"

| Mutant Construction | Forward primer | Reverse primer | Description |
|---|---|---|---|
| **Δ*lacI::kanR*** | TCTGGTGGCCGGAAGGCGAAGCGGCATGCATTTACGTTGAGTGTAGGCTGGAGCTGCTTC | TGCCTAATGAGTGAGCTAACTCACATTAATTGCGTTGCGCCATATGAATATCCTCCTTAG | Used to amplify *kanR* with FRT sites from pKD13, and the linear DNA was used with lambda Red system to replace *lacI* on the chromosome. |
| **Δ*araBAD*::$P_{T5}$-*gfp-kanR*** | GCCATAGCATTTTTATCCATAAGATTAGCGGATCCTACCTCTCGAGAAATCATAAAAAAT | AGCCTGGTTTCGTTTGATTGGCTGTGGTTTTATACAGTCATCAGAAGAACTCGTCAAGAA | Used to amplify $P_{T5}$-*gfp-kanR* from WM020, and the linear DNA was used with lambda Red system to replace *araBAD* on the chromosome. |
| **Δ*araBAD*::$P_{T5}$-*mcherry-kanR*** | GCCATAGCATTTTTATCCATAAGATTAGCGGATCCTACCTCTCGAGAAATCATAAAAAAT | AGCCTGGTTTCGTTTGATTGGCTGTGGTTTTATACAGTCATCAGAAGAACTCGTCAAGAA | Used to amplify $P_{T5}$-*mcherry-kanR* from MO001, and the linear DNA was used with lambda Red system to replace araBAD on the chromosome. |

**Table S2A. Primer sequences for mutant construction**

| Mutant Verification | External Forward | External Reverse | Internal Forward | Internal Reverse | Description |
|---|---|---|---|---|---|
| **Δ*hmp*** | TTTATAGGCGGTCCTGTTGG | AGGCGACCTTTTCAAGGAAT | TAACCGCCCATTTCTACGAC | TACGATAGCCTTTGCCATCC | *Absence of locus was confirmed by cPCR using the following combinations of primers:*<br>*1) External Forward, External Reverse*<br>*2) Internal Forward, Internal Reverse*<br>*3) External Forward, kanR Internal Reverse* |
| **Δ*greA*** | GTACCCGGGTCGGTATTTTT | CCACGCTCTGTTCGTTGATA | CCACGCTCTGTTCGTTGATA | ACAGCTTGGCTTCGATGTCT | *Absence of locus was confirmed by cPCR using the following combinations of primers:*<br>*1) External Forward, External Reverse*<br>*2) Internal Forward, Internal Reverse;*<br>*3) External Forward, kanR Internal Reverse* |
| **Δ*greB*** | CTCACGTTCGTCACCAGCTA | CATCGGCAGGAGGTTAAGAC | AAAAAGCGTCTGCGTGAAAT | TCGGGGAATCGATAGAGATG | *Absence of locus was confirmed by cPCR using the following combinations of primers:*<br>*1) External Forward, External Reverse*<br>*2) Internal Forward, Internal Reverse*<br>*3) External Forward, kanR Internal Reverse* |
| **Δ*lacI*** | GGGATCAGGAGGAGAAGATCG | | ATCGAATGGCGCAAAACCT | TTCCAGTCGGGAAACCTGT | *Absence of locus was confirmed by cPCR using the following combinations of primers:*<br>*1) Internal Forward, Internal Reverse*<br>*2) External Forward, kanR Internal Reverse* |
| **Δ*araBAD*::$P_{T5}$-*gfp/mcherry-kanR*** | | TCAGCCAGACAAACGTTGAG | ATGATTGAACAAGATGGATTGC | | *Absence of locus was confirmed by cPCR using the following combinations of primers:*<br>*1) Internal Forward, Internal Reverse* |
| ***kanR*** | | | | GAAGCGGTCAGCCCATTC | |

**Table S2B. Primer sequences for mutant verification**

| Plasmid Construction | Genotype | Forward primer | Reverse primer | Description |
|---|---|---|---|---|
| **pDS01** | pUA66 $P_{greA}$-*greA* | GAGGGGATCCACGATGACCCTTCGGGAACT | CATGCCTGCAGGGGTAATTCTTACAGGTATTC | Amplify $P_{greA}$-*greA* from MG1655 genomic DNA. Digest pUA66 and amplicon with BamHI and SbfI. Perform ligation with the Quick Ligation Kit (NEB). |
| **pDS02** | pUA66 $P_{greB}$-*greB* | GAGGGGATCCCAGCTTAAGTATACAATTTA | CATGCCTGCAGGTCTTCCTTACGGTTTCACGT | Amplify $P_{greB}$-*greB* from MG1655 genomic DNA. Digest pUA66 and amplicon with BamHI and SbfI. Perform ligation with the Quick Ligation Kit (NEB). |
| **pDS03** | puA66 $P_{T7}$-*gfp*, *lacIq* | GGGTTTTTTGCCAAGCTAGCTTGGCGAG | AGTCGTATTATTTCTCGAGCGATGACGTC | Amplify pSA21 with Phusion to be used in Hifi DNA assembly reaction. Double stranded DNA insert ($P_{T7}$-*gfp*) was purchased from Integrated DNA Technologies. |
| **pDS04** | pUA66 J23114 – *$T_7$RNAP*, $P_{T7}$-*gfp*, *lacIq* | CAAGAAAAGCCCGTCACGGGCTTC | TATTATGGGTAGTTTCCTTGCATGAATCCA | Amplify pDS03 with Phusion to be used in Hifi DNA assembly reaction. Double stranded DNA insert (J23114-*$T_7$RNAP*) was purchased from Integrated DNA Technologies. |
| **pDS05** | pQE80 $P_{T5}$-tdBroccoli, *lacIq* | CCTATTTCCCTAAAGGGTTTATTGAGAATATGTTTTTC | TTGAATCTATTATAATTGTTATCCGCTCACAAAGCAAATAAATTT | Amplify pXW09 with Phusion to be used in Hifi DNA assembly reaction. Double stranded DNA insert (tdBroccoli) was purchased from Integrated DNA Technologies. |
| **pDS06** | pQE80 $P_{T5}$-*mcherry*-tdBroccoli , *lacIq* | CTCTACGACAACCTCTTCACAGCCAATCTCGCCCGGATAGCTCAGTC | GCAAATAAATTTTTTATGATTTCTCGAG | Amplify pDS05 with Phusion to be used in Hifi DNA assembly reaction with *mcherry*. |
| | | CTCGAGAAATCATAAAAAAATTTATTTGC | GAGATTGGCTGTGAAGAGGTTGTCGTAGAGCTACTTGTACAGCTCGTCCA | Amplify *mcherry* from MO002 genomic DNA |
| **pDS07** | pUA66 $P_{hmp}$-*gfpsf*, $P_{greA}$-*greA* | CAAGAAAAGCCCGTCACGGGCTTC | TATTATGGGTAGTTTCCTTGCATGAATCCA | Amplify pXW02 with Phusion to be used in Hifi DNA assembly reaction. Double stranded DNA insert ($P_{greA}$-*greA*) was purchased from Integrated DNA Technologies. |
| **pDS08** | pUA66 $P_{hmp}$-*gfpsf*, $P_{greB}$-*greB* | CAAGAAAAGCCCGTCACGGGCTTC | TATTATGGGTAGTTTCCTTGCATGAATCCA | Amplify pXW02 with Phusion to be used in Hifi DNA assembly reaction. Double stranded DNA insert ($P_{greB}$-*greB*) was purchased from Integrated DNA Technologies. |

**Table S2C. Primer sequences for plasmid construction**

| Insert | Sequence |
|---|---|
| **pDS03 insert** | TAATACGACTCACTATAGATTCAATTGTGAGCGGATAACAATTTCACACAGAATTCATTAAAGAGGAGACCTCTAGTATGAGCAAAGGAGAAGAACTTTTCACTGGAGTTGTCCCAATTCTTGTTGAATTAGATGGTGATGTTAATGGGCACAAATTTTCTGTCCGTGGAGAGGGTGAAGGTGATGCTACAAACGGAAAACTCACCCTTAAATTTATTTGCACTACTGGAAAACTACCTGTTCCGTGGCCAACACTTGTCACTACTCTGACCTATGGTGTTCAATGCTTTTCCCGTTATCCGGATCACATGAAACGGCATGACTTTTTCAAGAGTGCCATGCCCGAAGGTTATGTACAGGAACGCACTATATCTTTCAAAGATGACGGGACCTACAAGACGCGTGCTGAAGTCAAGTTTGAAGGTGATACCCTTGTTAATCGTATCGAGTTAAAGGGTATTGATTTTAAAGAAGATGGAAACATTCTTGGACACAAACTCGAGTACAACTTTAACTCACACAATGTATACATCACGGCAGACAAACAAAAGAATGGAATCAAAGCTAACTTCAAAATTCGCCACAACGTTGAAGATGGTTCCGTTCAACTAGCAGACCATTATCAACAAAATACTCCAATTGGCGATGGCCCTGTCCTTTTACCAGACAACCATTACCTGTCGACACAATCTGTCCTTTCGAAAGATCCCAACGAAAAGCGTGACCACATGGTCCTTCTTGAGTTTGTAACTGCTGCTGGGATTACACATGGCATGGATGAGCTCTACAAATAAGGATCCGCATGCGAGCTCGGTACCCCGGGTCGACCTGCAGCCAAGCTTAATTAGCTGAGCTTGTAGCATAACCCCTTGGGGCCTCTAAACGGGTCTTGAGGGGTTTTTTG |
| **pDS04 insert** | CAAGGAAACTACCCATAATAtttatggctagctcagtcctaggtacaatgctagcTTCACACAGAATTCATTAAAGAGGAGAAATTAACTatgAACACGATTAACATCGCTAAGAACGACTTCTCTGACATCGAACTGGCTGCTATCCCGTTCAACACTCTGGCTGACCATTACGGTGAGCGTTTAGCTCGCGAACAGTTGGCCCTTGAGCATGAGTCTTACGAGATGGGTGAAGCACGCTTCCGCAAGATGTTTGAGCGTCAACTTAAAGCTGGTGAGGTTGCGGATAACGCTGCCGCCAAGCCTCTCATCACTACCCTACTCCCTAAGATGATTGCACGCATCAACGACTGGTTTGAGGAAGTGAAAGCTAAGCGCGGCAAGCGCCCGACAGCCTTCCAGTTCCTGCAAGAAATCAAGCCGGAAGCCGTAGCGTACATCACCATTAAGACCACTCTGGCTTGCCTAACCAGTGCTGACAATACAACCGTTCAGGCTGTAGCAAGCGCAATCGGTCGGGCCATTGAGGACGAGGCTCGCTTCGGTCGTATCCGTGACCTTGAAGCTAAGCACTTCAAGAAAAACGTTGAGGAACAACTCAACAAGCGCGTAGGGCACGTCTACAAGAAAGCATTTATGCAAGTTGTCGAGGCTGACATGCTCTCTAAGGGTCTACTCGGTGGCGAGGCGTGGTCTTCGTGGCATAAGGAAGACTCTATTCATGTAGGAGTACGCTGCATCGAGATGCTCATTGAGTCAACCGGAATGGTTAGCTTACACCGCCAAAATGCTGGCGTAGTAGGTCAAGACTCTGAGACTATCGAACTCGCACCTGAATACGCTGAGGCTATCGCAACCCGTGCAGGTGCGCTGGCTGGCATCTCTCCGATGTTCCAACCTTGCGTAGTTCCTCCTAAGCCGTGGACTGGCATTACTGGTGGTGGCTATTGGGCTAACGGTCGTCGTCCTCTGGCGCTGGTGCGTACTCACAGTAAGAAAGCACTGATGCGCTACGAAGACGTTTACATGCCTGAGGTGTACAAAGCGATTAACATTGCGCAAAACACCGCATGGAAAATCAACAAGAAAGTCCTAGCGGTCGCCAACGTAATCACCAAGTGGAAGCATTGTCCGGTCGAGGACATCCCTGCGATTGAGCGTGAAGAACTCCCGATGAAACCGGAAGACATCGACATGAATCCTGAGGCTCTCACCGCGTGGAAACGTGCTGCCGCTGCTGTGTACCGCAAGGACAAGGCTCGCAAGTCTCGCCGTATCAGCCTTGAGTTCATGCTTGAGCAAGCCAATAAGTTTGCTAACCATAAGGCCATCTGGTTCCCTTACAACATGGACTGGCGCGGTCGTGTTTACGCTGTGTCAATGTTCAACCCGCAAGGTAACGATATGACCAAAGGACTGCTTACGCTGGCGAAAGGTAAACCAATCGGTAAGGAAGGTTACTACTGGCTGAAAATCCACGGTGCAAACTGTGCGGGTGTCGATAAGGTTCCGTTCCCTGAGCGCATCAAGTTCATTGAGGAAAACCACGAGAACATCATGGCTTGCGCTAAGTCTCCACTGGAGAACACTTGGTGGGCTGAGCAAGATTCTCCGTTCTGCTTCCTTGCGTTCTGCTTTGAGTACGCTGGGGTACAGCACCACGGCCTGAGCTATAACTGCTCCCTTCCGCTGGCGTTTGACGGGTCTTGCTCTGGCATCCAGCACTTCTCCGCGATGCTCCGAGATGAGGTAGGTGGTCGCGCGGTTAACTTGCTTCCTAGTGAAACCGTTCAGGACATCTACGGGATTGTTGCTAAGAAAGTCAACGAGATTCTACAAGCAGACGCAATCAATGGGACCGATAACGAAGTAGTTACCGTGACCGATGAGAACACTGGTGAAATCTCTGAGAAAGTCAAGCTGGGCACTAAGGCACTGGCTGGTCAATGGCTGGCTTACGGTGTTACTCGCAGTGTGACTAAGCGTTCAGTCATGACGCTGGCTTACGGGTCCAAAGAGTTCGGCTTCCGTCAACAAGTGCTGGAAGATACCATTCAGCCAGCTATTGATTCCGGCAAGGGTCTGATGTTCACTCAGCCGAATCAGGCTGCTGGATACATGGCTAAGCTGATTTGGGAATCTGTGAGCGTGACGGTGGTAGCTGCGGTTGAAGCAATGAACTGGCTTAAGTCTGCTGCTAAGCTGCTGGCTGCTGAGGTCAAAGATAAGAAGACTGGAGAGATTCTTCGCAAGCGTTGCGCTGTGCATTGGGTAACTCCTGATGGTTTCCCTGTGTGGCAGGAATACAAGAAGCCTATTCAGACGCGCTTGAACCTGATGTTCCTCGGTCAGTTCCGCTTACAGCCTACCATTAACACCAACAAAGATAGCGAGATTGATGCACACAAACAGGAGTCTGGTATCGCTCCTAACTTTGTACACAGCCAAGACGGTAGCCACCTTCGTAAGACTGTAGTGTGGGCACACGAGAAGTACGGAATCGAATCTTTTGCACTGATTCACGACTCCTTCGGTACCATTCCGGCTGACGCTGCGAACCTGTTCAAAGCAGTGCGCGAAACTATGGTTGACACATATGAGTCTTGTGATGTACTGGCTGATTTCTACGACCAGTTCGCTGACCAGTTGCACGAGTCTCAATTGGACAAAATGCCAGCACTTCCGGCTAAAGGTAACTTGAACCTCCGTGACATCTTAGAGTCGGACTTCGCGTTCGCGtaaGGATCCGCATGCGAGCTCGGTACCCCGGGTCGACCTGCAGCCAAGCTTAATTAGCTGAGCTTGGACTCCTGTTGATAGATCCAGTAATGACCTCAGAACTCCATCTGGATTTGTTCAGAACGCTCGGTTGCCGCCGGGCGTTTTTTATTGGTGAGAATCAAGAAAAGCCCGTCACGGG |
| **pDS05 insert** | AACAATTATAATAGATTCAATTGTGAGCGGATAACAATTTCGCCCGGATAGCTCAGTCGGTAGAGCAGCGGAGACGGTCGGGTCCATCTGAGACGGTCGGGTCCAGATATTCGTATCTGTCGAGTAGAGTGTGGGCTCAGATGTCGAGTAGAGTGTGGGCTCCGCGGGTCCAGGGTTCAAGTCCCTGTTCGGGCGCCAGACTCCTGTTGATAGATCCAGTAATGACCTCAGAACTCCATCTGGATTTGTTCAGAACGCTCGGTTGCCGCCGGGCGTTTTTTATTGGTGAGAATCCAAGCTAGCTTGGCGAGATTTTCAGGAGCTAAGGAAGCTAAAATGGAGAAAAAAATCACTGGATATACCACCGTTGATATATCCCAATGGCATCGTAAAGAACATTTTGAGGCATTTCAGTCAGTTGCTCAATGTACCTATAACCAGACCGTTCAGCTGGATATTACGGCCTTTTTAAAGACCGTAAAGAAAAATAAGCACAAGTTTTATCCGGCCTTTATTCACATTCTTGCCCGCCTGATGAATGCTCATCCGGAATTTCGTATGGCAATGAAAGACGGTGAGCTGGTGATATGGGATAGTGTTCACCCTTGTTACACCGTTTTCCATGAGCAAACTGAAACGTTTTCATCGCTCTGGAGTGAATACCACGACGATTTCCGGCAGTTTCTACACATATATTCGCAAGATGTGGCGTGTTACGGTGAAAACCTGGCCTATTTCCCTAAAGGGTTT |
| **pDS07 insert** | CAAGGAAACTACCCATAATAGATCCACGATGACCCTTCGGGAACTTCAGGGTAAAATGACTATCAAAATGTGAATTGTAGCTGACCTGGGACTTGTACCCGGGTCGGTATTTTTTTGCTTCTGGTCCCGGTAAGGAGTTATGCCGGGCAGGCCGAACAGCCGGGGTGGGTGAAGACTTGCCCTATCAGGAATATTCAAGAGGTATAACAAatgCAAGCTATTCCGATGACCTTACGCGGCGCTGAAAAATTACGCGAAGAGCTGGATTTTCTGAAATCTGTGCGCCGTCCTGAAATCATTGCTGCTATCGCGGAAGCGCGTGAGCATGGCGACCTGAAAGAAAACGCCGAATACCACGCAGCTCGTGAACAGCAGGGTTTCTGCGAAGGCCGTATTAAAGACATCGAAGCCAAGCTGTCGAACGCGCAGGTGATTGATGTCACCAAAATGCCCAACAATGGGCGCGTTATTTTTGGTGCTACCGTAACGGTGCTGAATCTGGATTCTGACGAAGAACAGACTTATCGCATCGTTGGCGATGACGAAGCTGACTTTAAACAAAACCTGATTTCTGTAAACTCGCCTATTGCTCGTGGCCTGATCGGCAAAGAAGAAGATGATGTTGTGGTCATCAAAACGCCGGGCGGCGAAGTAGAATTTGAAGTAATTAAGGTGGAATACCTGtaaGAATTACCCCTGCAGGCATGCAAGCTCTAGAGGCATCAAATAAAACGAAAGGCTCAGTCGAAAGACTGGGCCTTTCGTTTTATCTGTTGTTTGTCGGTGAACGCTCTCCTGAGTAGGACAAATCAAGAAAAGCCCGTCACGGG |
| **pDS08 insert** | CAAGGAAACTACCCATAATAGATCCCAGCTTAAGTATACAATTTATTCGGCGAAACATTATTGATTCTGTTGATATGATCACGTTATACCCAATGTGCGCATTATCAAACAGACAAAGGGAATCAACGAGatgAAAACGCCCCTGGTTACCCGGGAAGGGTATGAAAAACTCAAACAAGAGCTTAATTATCTCTGGCGTGAAGAACGCCCGGAGGTCACAAAAAAGGTGACCTGGGCCGCAAGTCTGGGCGACCGCAGCGAAAATGCTGACTATCAGTATAATAAAAAGCGTCTGCGTGAAATCGACCGTCGCGTGCGCTATCTCACTAAATGCCTGGAAAATCTCAAAATCGTCGATTACTCCCCTCAGCAGGAAGGCAAAGTCTTTTTTGGCGCGTGGGTGGAGATTGAAAACGACGATGGCGTGACTCACCGTTTCCGTATTGTCGGCTACGATGAAATTTTTGGCCGTAAAGATTACATCTCTATCGATTCCCCGATGGCCCGCGCATTGCTGAAAAAAGAAGTCGGCGATCTGGCGGTGGTGAATACCCCTGCCGGGGAAGCGAGCTGGTATGTTAATGCTATCGAGTACGTGAAACCGtaaGGAAGACCTGCAGGCATGCAAGCTCTAGAGGCATCAAATAAAACGAAAGGCTCAGTCGAAAGACTGGGCCTTTCGTTTTATCTGTTGTTTGTCGGTGAACGCTCTCCTGAGTAGGACAAATCAAGAAAAGCCCGTCACGGG |

**Table S2D. Double stranded insert sequences**
